# Regulation of REM Sleep by Inhibitory Neurons in the Dorsomedial Medulla

**DOI:** 10.1101/2020.11.30.405530

**Authors:** Joseph A. Stucynski, Amanda L. Schott, Justin Baik, Shinjae Chung, Franz Weber

## Abstract

The two major stages of mammalian sleep – rapid eye movement sleep (REMs) and non-REM sleep (NREMs) – are characterized by distinct brain rhythms ranging from millisecond to minute-long (infraslow) oscillations. The mechanisms controlling transitions between sleep stages and how they are synchronized with infraslow rhythms remain poorly understood. Using opto- and chemogenetic manipulation, we show that GABAergic neurons in the dorsomedial medulla (dmM) promote the initiation and maintenance of REMs, in part through their projections to the dorsal and median raphe nuclei. Fiber photometry revealed that dmM GABAergic neurons are strongly activated during REMs. During NREMs, their activity fluctuated in close synchrony with infraslow oscillations in the spindle band of the electroencephalogram, and the phase of this rhythm modulated the latency of optogenetically induced REMs episodes. Thus, dmM inhibitory neurons powerfully promote REMs, and their slow activity fluctuations may coordinate transitions from NREMs to REMs with infraslow brain rhythms.

## INTRODUCTION

Mammalian sleep comprises two major substages: rapid eye movement sleep (REMs) and non-REM sleep (NREMs). Both stages are characterized by distinct oscillations in the electroencephalogram (EEG) that typically reside in the sub-second range, such as slow waves, sleep spindles, and hippocampal theta oscillations (Adamantidis et al., 2019; Saper et al., 2010; Scammell et al., 2017). In addition to these fast oscillations, the sleep architecture is also shaped by much slower processes such as the ultradian NREMs/REMs cycle or infraslow rhythms operating on a minute timescale (Fernandez and Lüthi, 2019; Lecci et al., 2017; Terzano et al., 1985; Watson, 2018). During NREMs, the EEG follows a distinct infraslow (∼0.02 Hz) oscillation in the sleep spindle (sigma) range, which plays a prominent role in timing transitions from NREMs to wakefulness (Lecci et al., 2017). However, the mechanisms by which infraslow processes interact with sleep circuits to coordinate brain state switches, particularly NREMs to REMs transitions, remain largely unknown.

Neural circuits involved in regulating transitions from NREMs to REMs have been identified in the hypothalamus and brainstem, including pons and medulla (Luppi et al., 2012; Park and Weber, 2020; Peever and Fuller, 2017; Saper et al., 2010; Scammell et al., 2017). Within the medulla, GABAergic neurons located in its ventral part have been shown to strongly promote REMs, partly through inhibition of REMs-suppressing (REMs-off) neurons in the ventrolateral periaqueductal gray (vlPAG) within the midbrain (Weber et al., 2015). In addition to the ventral medulla, the dorsomedial medulla (dmM) has also been implicated in REMs control (Clément et al., 2012; Clément et al., 2014, Gervasoni et al., 2000; Goutagny et al., 2008; Kaur et al., 2001; Sakai, 2018; Sapin et al., 2009; Verret et al., 2006). Electrophysiological *in vivo* recordings in rodents demonstrated the existence of REMs-active neurons in both the nucleus prepositus hypoglossi (PH) and the dorsal paragigantocellular reticular nucleus (DPGi) (Goutagny et al., 2008; Sakai, 2018), and pharmacological inhibition of DPGi neurons expressing adrenergic alpha2-receptors specifically reduced the amount of REMs (Clément et al., 2014). Inhibitory neurons in the PH and DPGi are thought to promote REMs by inhibiting REMs-off neurons in the pons and midbrain, located within the locus coeruleus (LC), dorsal raphe (DRN), and vlPAG (Ennis and Aston-Jones, 1989; Gervasoni et al., 2000; Goutagny et al., 2008; Clément et al., 2012; Sapin et al., 2009; Verret et al., 2006). DPGi GABAergic neurons, and neurons within the DPGI that project to the vlPAG or LC, express high levels of the immediate early gene c-Fos after deprivation-induced REMs rebound, suggesting a strong activation during REMs (Clément et al., 2012; Sapin et al., 2009; Verret et al., 2006). The slow time course of c-Fos expression, however, lacks the temporal precision to resolve activity changes associated with fast brain state transitions. Moreover, electrical stimulation of the PH resulted in an increased amount of REMs, an effect which could be reversed by pharmacological inhibition of the LC (Kaur et al., 2001), but non-specific activation of axons-of-passage causing this effect cannot be ruled out. Although these studies suggest an important role of dmM inhibitory neurons in REMs regulation, their precise role in initiating and maintaining REMs and the underlying circuit mechanisms are still not fully understood. Furthermore, since the oscillation in EEG sigma power influences the timing of awakenings from NREMs (Lecci et al., 2017), an interesting question is whether this infraslow rhythm is also involved in timing transitions from NREMs to REMs, possibly by influencing the activity of REMs-regulatory circuits such as the dmM inhibitory neurons.

In this study, we found that optogenetic activation of GABAergic, GAD2-expressing neurons in the dmM powerfully promoted the initiation and maintenance of REMs, while chemogenetic inactivation reduced the amount of REMs. The REMs-promoting effect was in part mediated by dmM neurons that project to the dorsal and median raphe (DR/MRN). Calcium imaging using fiber photometry revealed that dmM neurons are strongly activated during REMs. During NREMs, their activity closely followed the infraslow oscillation in the EEG sigma power. In addition, we found that this oscillation modulated the latency of optogenetically induced REMs episodes. Our findings delineate the role of dmM inhibitory neurons in REMs control and suggest a mechanism by which infraslow oscillations contribute to the timing of NREMs to REMs transitions.

## RESULTS

### Optogenetic activation of dmM GAD2 neurons powerfully promotes REMs

To probe the role of dmM GABAergic neurons in sleep-wake control, we injected Cre-inducible adeno-associated viruses (AAVs) expressing channelrhodopsin-2 fused with enhanced yellow fluorescent protein (AAV-DIO-ChR2-eYFP) into the dmM of GAD2-Cre mice (**Fig. 1a**). ChR2-eYFP was consistently expressed in the PH and DPGi across mice and to a lesser extent in neighboring areas including the medial vestibular nucleus (MV) and nucleus of the solitary tract (NST) (**Fig. 1a**,**b**). The optic fibers for laser stimulation were consistently placed on top of the PH (**Suppl. Fig. 1a**). While monitoring the animal’s brain state using electroencephalogram (EEG) and electromyogram (EMG) recordings, laser stimulation (10 Hz, 120 s per trial) was applied randomly every 10 to 20 minutes (**Fig. 1c; Suppl. Video 1**). We found that optogenetic activation of dmM GAD2 neurons strongly increased the percentage of REMs during the laser stimulation interval (P < 0.001, bootstrap, n = 9 mice), while NREMs was reduced (**Fig. 1d**; P < 0.001). The percentage of wakefulness was also enhanced (P < 0.001), but increased at a slower rate than REMs. In control mice expressing eYFP only, laser stimulation did not significantly alter the brain state (**Suppl. Fig. 1b**; P > 0.233, n = 7 mice). Consistent with the strong REMs-promoting effect, the mean EEG spectrogram averaged across laser trials displayed a distinct increase in both theta and gamma power with a concomitant power reduction in the delta and sigma ranges during laser stimulation (**Fig. 1e**; delta, P = 3.5e-5, T = −8.26; theta, P = 0.007, T = 3.60; sigma, P = 0.001, T = −4.91; gamma, P = 6.8e-5, T = 7.52; paired t-test, n = 9 mice).

**Figure 1.**
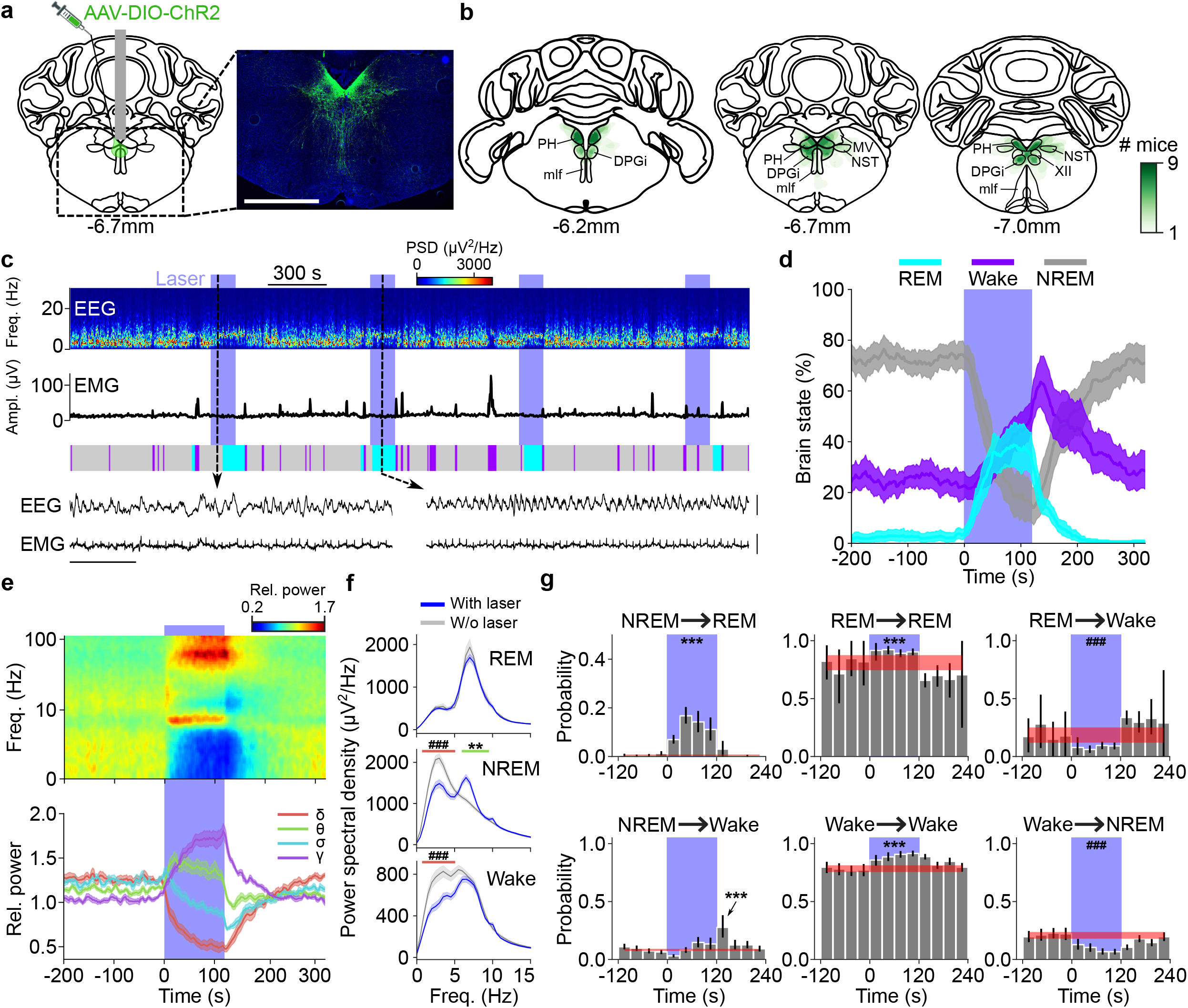
Optogenetic activation of dmM GAD2 neurons powerfully promotes REMs. **(a)** Schematic of optogenetic experiment. Left, coronal diagram of mouse brain indicating injection site and optic fiber placement. Gray rectangle, optic fiber. Right, fluorescence image of dmM in a GAD2-Cre mouse injected with ChR2-eYFP (green). Blue, Hoechst stain. Scale bar, 1 mm. **(b)** Outline of areas with ChR2-eYFP expressing cell bodies along the rostrocaudal axis within three coronal brain sections. The green color code indicates in how many mice the virus expression overlapped at the corresponding location (n = 9 mice). The coronal brain schemes were adapted from Allen Mouse Brain Atlas (© 2015 Allen Institute for Brain Science. Allen Brain Atlas API. Available from: http://brain-map.org/api/index.html). PH, nucleus prepositus hypoglossi; DPGi, dorsal paragigantocellular nucleus; mlf, medial longitudinal fasciculus; MV, medial vestibular nucleus; NTS, nucleus of the solitary tract; XII, hypoglossal nucleus. **(c)** Example experiment. Shown are EEG spectrogram, EMG amplitude, and color-coded brain states. The blue patches indicate 2 min laser stimulation intervals (473 nm, 10 Hz). Two EEG and EMG raw traces are shown at an expanded timescale for the selected time points (dashed lines; scale bars, 1 s and 0.5 mV). PSD, power spectral density. **(d)** Percentage of REMs, NREMs, and wake before, during, and after laser stimulation (n = 9 mice). Laser stimulation significantly increased REMs (P < 0.001, bootstrap, n = 9 mice), decreased NREMs (P < 0.001) and caused a delayed increase in wake (P < 0.001). Shadings, 95% confidence intervals (CIs). **(e)** Impact of laser stimulation on EEG spectrogram and different power bands. Top, laser trial averaged EEG spectrogram with logarithmic frequency axis. Each frequency component of the spectrogram was normalized by its mean power across the recording. Bottom, time course of delta (0.5 − 4.5 Hz), theta (6 − 9 Hz), sigma (10 − 15 Hz), and gamma power (55 − 90 Hz) before, during, and after laser stimulation. During laser stimulation, theta (P = 0.007, T = 3.60; paired t-test, n = 9 mice) and gamma power (P = 6.8e-5, T = 7.52) were increased, while delta (P = 3.5e-5, T = −8.26) and sigma power (P = 0.001, T = −4.91) were reduced. Shadings, ± s.e.m. **(f)** Power spectral density of EEG during REMs (top), NREMs (middle), and wake (bottom) with and without (w/o) laser stimulation. During NREMs, laser stimulation significantly reduced delta power (red line; P = 0.00025, T = −6.22, paired t-test) and increased theta power (green line; P = 9.9e-4, T = 5.04). During wake, the delta power (P = 0.0001, T = −6.88) was reduced. **/^##^, P < 0.01; ***/^###^, P < 0.001 for significant increases/decreases. **(g)** Effect of laser stimulation of dmM GAD2 neurons on brain state transition probabilities. Transition probabilities were calculated for 10 s bins. Each bar represents the average transition probability during a 30 s interval. Error bars represent the 95% CI. The red line and shading depict the average baseline transition probability (computed for the interval preceding laser stimulation) and the 95% CI, respectively. Laser stimulation significantly increased the probability of NREMs to REMs transitions (NREM → REM, P < 0.001, bootstrap), REMs to REMs (REM → REM, P < 0.001) and wake to wake transitions (Wake → Wake, P < 0.001), while suppressing REMs to wake (REM → Wake, P < 0.001) and wake to NREMs transitions (Wake → NREM, P < 0.001). The arrow indicates a brief increase in NREMs to wake transitions directly following laser stimulation (P < 0.001). CIs and P-values were calculated using bootstrapping. ***/^###^, P < 0.001 for significant increases/decreases.

The effect of optogenetically activating dmM GAD2 neurons on the EEG was dependent on the brain state. The power spectral density of the EEG during laser-induced REMs episodes was indistinguishable from that during spontaneous REMs, without significant changes in the delta, theta, or sigma power (**Fig. 1f;** REM, P > 0.48 for delta, theta, and sigma; paired t-test, n = 9 mice; **Suppl. Fig. 1c**) and the EMG amplitude was also unchanged (**Suppl. Fig. 1e**; P = 0.11, paired t-test). In contrast, laser stimulation strongly reduced the EEG delta power during both NREMs and wakefulness and increased the NREMs theta power (**Fig. 1f**; NREM, delta, P = 0.00025, T = −6.22; theta, P = 0.001, T = 5.04; Wake, delta, P = 0.0001, T = −6.88; paired t-test). Activation of dmM neurons during NREMs thus caused changes in the EEG resembling two defining features of REMs – reduced delta and increased theta power – which also precede spontaneous NREMs to REMs transitions (Gottesmann, 1996) (**Suppl. Fig. 1c**).

Next, to determine whether the changes in the percentage of a specific brain state during laser stimulation are due to changes in the induction or maintenance of that state, we quantified how the laser affected the transition probability between each pair of brain states. Activation of dmM neurons caused an increase in NREMs to REMs transitions (**Fig. 1g**; NREM → REM, P < 0.001, bootstrap, n = 9 mice), enhanced REMs to REMs transitions (REM → REM, P < 0.001), and suppressed REMs to wake transitions (REM → Wake, P < 0.001), suggesting that dmM GAD2 neurons promote both the initiation and maintenance of REMs. Interestingly, in rare instances laser stimulation also promoted transitions from wakefulness to REMs (Wake → REM, P < 0.001). In each of these cases, the wake period interrupted two successive REMs episodes (**Suppl. Fig. 1d**,**f**).

In addition, optogenetic activation also promoted wake to wake transitions (Wake → Wake, P < 0.001), while the probability of NREMs to wake transitions was unchanged (**Fig. 1g**; NREM → Wake, P = 0.27). Hence, the increase of the wake percentage during laser stimulation was due to an enhanced maintenance (instead of induction) of wakefulness. Immediately after cessation of laser stimulation, the probability of NREMs to wake transitions was briefly increased (NREM → Wake, P < 0.001, bootstrap), possibly due to a rebound in the activity of wake-promoting neurons such as adrenergic/noradrenergic neurons, which have been shown to be inhibited by the dmM (Clément et al., 2014; Verret et al., 2006; Kaur et al., 2001; Ennis and Aston-Jones, 1989). Taken together, our optogenetic activation experiments therefore suggest that dmM GAD2 neurons both initiate and maintain REMs and are also involved in the maintenance of wakefulness.

### Closed-loop optogenetic activation of dmM GAD2 neurons maintains REMs

The transition analysis indicated that activation of dmM GAD2 neurons prolongs REMs (**Fig. 1g;** REM → REM). To more directly test the role of these neurons in REMs maintenance, we applied optogenetic closed-loop stimulation (**Fig. 2a**,**b**). The brain state of the animal was classified based on real-time analysis of the EEG/EMG signals. As soon as the onset of a spontaneous REMs episode was detected, laser stimulation was initiated, and stayed on until the episode ended. The laser was turned on randomly for only 50% of the detected REMs episodes. Optogenetic closed-loop activation of dmM neurons significantly prolonged the duration of REMs episodes overlapping with laser stimulation (**Fig. 2c**; no laser, 63.5 s; laser, 118.5 s; P = 0.0003, T = 11.70, paired t-test, n = 5 mice). In contrast, closed-loop stimulation in eYFP control mice had no significant effect on the REMs duration (**Suppl. Fig. 2a;** no laser, 86.7 s; laser, 88.3 s; P = 0.80, T = 0.27, paired t-test, n = 6 mice). These results further demonstrate a prominent role of dmM GAD2 neurons in powerfully maintaining REMs.

**Figure 2.**
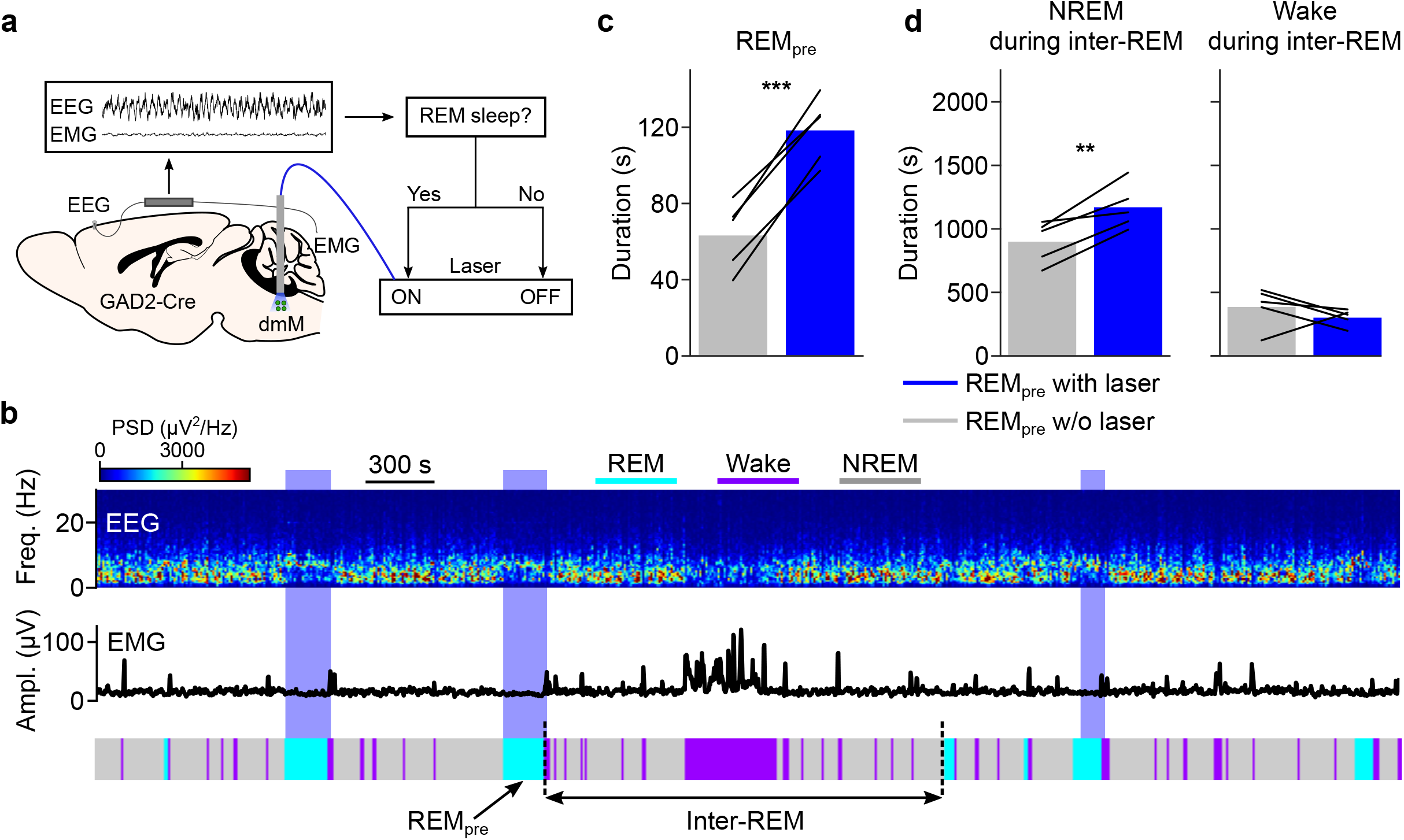
Closed-loop optogenetic activation of dmM GAD2 neurons maintains REMs. **(a)** Schematic of closed-loop stimulation protocol. The brain state was continuously monitored; once a REMs episode was detected, laser stimulation was initiated and maintained throughout REMs. **(b)** Example experiment. Shown are EEG spectrogram, EMG amplitude, and color-coded brain states. The blue patches indicate laser stimulation intervals (473 nm, 10 Hz). The arrows indicate a preceding REMs episode (REM_pre_) and the subsequent inter-REM interval. PSD, power spectral density. **(c)** Duration of REMs episodes with and without laser. Closed-loop stimulation significantly increased the duration of REMs episodes (P = 0.0003, T = 11.70, paired t-test, n = 5 mice). ***, P < 0.001. Bars, mean duration; lines, individual mice. **(d)** Effects of laser stimulation on the total NREMs and wake duration during the inter-REM interval. REMs episodes with closed-loop stimulation were followed by a larger total duration of NREMs during the subsequent inter-REM interval (NREM, P = 0.009, T = 4.71; paired t-test, n = 5 mice). The subsequent amount of wakefulness was not significantly changed (Wake, P = 0.35, T = −1.06). *, P < 0.05. Bars, mean duration; lines, individual mice.

As shown in multiple mammalian species, the interval between two successive REMs periods (inter-REM interval) is positively correlated with the preceding REMs duration, mainly due to a larger amount of NREMs following longer REMs episodes (Barbato and Wehr, 1998; Benington and Heller, 1994; Ursin, 1970; Vivaldi et al., 1994). This correlation provides evidence for a homeostatic process regulating the timing of REMs episodes on the ultradian timescale (Benington and Heller, 1994; Le Bon, 2020; Ocampo-Garcés et al., 2000, 2020), in which homeostatic pressure for REMs building up during the inter-REM interval is dissipated during each REMs episode in proportion to its duration. As further support for the homeostatic regulation of REMs, we found that REMs periods overlapping with laser stimulation were followed by a larger amount of NREMs during the subsequent inter-REM interval, while the amount of wakefulness was not significantly altered (**Fig. 2d**; NREM, P = 0.009, T = 4.71; Wake, P = 0.350, T = −1.06, paired t-test, n = 5 mice). This finding suggests that more REMs pressure is discharged during REMs periods extended by laser stimulation, requiring a longer duration of subsequent NREMs to accumulate sufficient pressure to re-enter REMs.

### Chemogenetic inhibition of dmM GAD2 neurons suppresses REMs

To test whether dmM GAD2 neurons also play a necessary role in REMs regulation, we chemogenetically inhibited their activity by injecting AAV8-hSyn-DIO-hM4D(Gi)-mCherry into the dmM of GAD2-Cre mice (**Fig. 3a**). hM4D(Gi)-mCherry was consistently expressed in the PH and DPGi across mice with only minor expression in neighboring areas (**Fig. 3b**). Compared to saline injection, clozapine-N-oxide (CNO) injection decreased the percentage of REMs during the first four hours in hM4D(Gi) mice (**Fig. 3c**,**d** top) and had no significant effect in control mice expressing mCherry (**Fig. 3d** bottom; mixed ANOVA with virus (mCherry or hm4D(Gi)) and intervention (saline or CNO) as between and within factors; interaction − virus * intervention, P = 1.4e-5, F(1, 14) = 42.41; Bonferroni correction, effect of CNO vs saline in hM4D(Gi) mice, P = 0.0011, T = 5.93; n = 8 hM4D(Gi) and n = 8 mCherry mice). The effect of CNO on REMs was also significantly different between the mCherry- and hM4D(Gi)-expressing mice (Bonferroni correction, effect of CNO in mCherry vs hm4Di mice, P = 0.0095, T = 3.35). Despite an increase in NREMs in hM4D(Gi) mice (interaction − virus * intervention, P = 0.049, F(1,14) = 4.66; Bonferroni correction, effect of CNO vs saline in hM4D(Gi) mice, P = 7.7e-4, T = 6.35), the ratio of REMs relative to total sleep (REMs / (NREMs + REMs)) was reduced (**Fig. 3e**; interaction − virus * intervention, P = 7.85e-07, F(1,14) = 70.37; Bonferroni correction, effect of CNO vs saline in hM4D(Gi) mice, P = 2.0e-6, T = 16.28). The decrease in REMs was due to a reduction in the frequency of REMs episodes (**Fig. 3f**) (interaction − virus*intervention, P = 0.0064, F(1,14) = 10.27; Bonferroni correction, effect of CNO vs saline in hm4D(Gi) mice, P = 0.012, T = 3.88). These results further emphasize the important role of dmM GAD2 neurons in REMs regulation.

**Figure 3.**
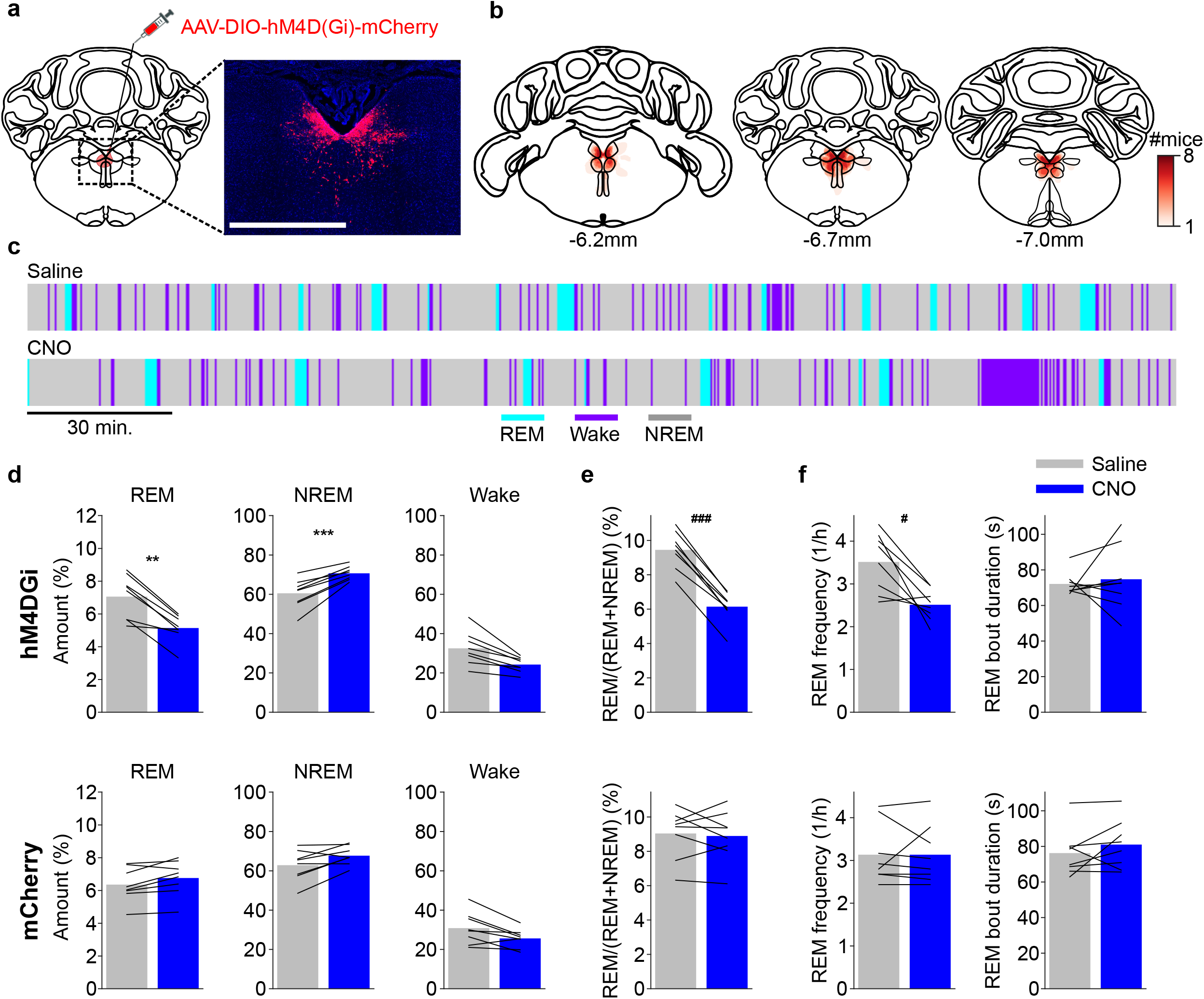
Chemogenetic inhibition of dmM GAD2 neurons suppresses REMs. **(a)** Schematic of chemogenetic inhibition experiment. Left, coronal diagram of mouse brain. Right, fluorescence image of dmM in a GAD2-Cre mouse injected with hM4D(Gi)-mCherry (red). Blue, Hoechst stain. Scale bar, 1 mm. **(b)** Heatmap outlining cell bodies expressing hM4D(Gi) within three consecutive sections along the rostrocaudal axis, n = 8 mice. **(c)** Example hypnograms of a 4 h control (saline) and CNO session from one mouse. The recording started 30 min after saline or CNO injection. **(d)** Impact of CNO in mice expressing hM4D(Gi) and mCherry. Top, CNO significantly reduced the percentage of REMs (mixed ANOVA with virus (mCherry or hm4D(Gi)) and intervention (saline or CNO) as between and within factors; interaction - virus * intervention, P = 1.4e-5, F(1, 14) = 42.41; Bonferroni correction, effect of CNO vs saline in hM4D(Gi) mice, P = 0.0011, T = 5.93; n = 8 hM4D(Gi) and n = 8 mCherry mice) and increased NREMs (interaction - virus * intervention, P = 0.049, F(1,14) = 4.66; Bonferroni correction, effect of CNO vs saline in hM4D(Gi) mice, P = 7.7e-4, T = 6.35). Bottom, effects of CNO in mCherry mice. **, P < 0.01; ***, P < 0.001 for significant increases. **(e)** CNO significantly reduced the ratio of REMs to total sleep (REMs / (REMs + NREMs)) in hM4D(Gi) mice (interaction - virus * intervention, P = 7.85e-07, F(1,14) = 70.37; Bonferroni correction, effect of CNO vs saline in hM4D(Gi) mice, P = 2.0e-6, T = 16.28). ^###^, P < 0.001 for significant decreases. **(f)** Effects of CNO on REMs frequency and duration. In hm4D(Gi) mice, CNO injection reduced the frequency of REMs episodes (interaction - virus * intervention, P = 0.0064, F(1,14) = 10.27; Bonferroni correction, effect of CNO vs saline in hM4D(Gi) mice, P = 0.012, T = 3.88). ^#^, P < 0.05 for significant decreases.

### DR/MRN-projecting dmM GAD2 neurons specifically promote REMs

Our optogenetic experiments showed that activation of dmM GAD2 neurons both promotes REMs and maintains wakefulness. We hypothesized that the effects on REMs and wakefulness may be mediated by different subpopulations within the dmM. Furthermore, we reasoned that the axons of REMs-promoting dmM neurons project to areas containing REMs-suppressing (REMs-off) neurons. Anterograde tracing using ChR2-eYFP revealed strong axonal projections of the dmM GAD2 neurons to the dorsal and median raphe nuclei (DR/MRN) at the midbrain/pons boundary and only sparse projections to the vlPAG (**Fig. 4a**). A recent study has shown that tonic optogenetic activation of serotonergic neurons in the DRN strongly suppresses REMs, while maintaining NREMs (Oikonomou et al., 2019); an effect comparable to that of stimulation of vlPAG GABAergic neurons, which form another prominent group of REMs-off neurons in the midbrain (Hayashi et al., 2015; Kaur et al., 2009; Lu et al., 2006; Sapin et al., 2009; Vanini et al., 2007; Weber et al., 2018). To specifically express ChR2-eYFP in the subpopulation of dmM GAD2 neurons projecting to the raphe nuclei we injected an AAV with high retrograde efficiency (AAVrg-DIO-ChR2-eYFP) into the DR/MRN (Tervo et al., 2016) (**Fig. 4b**). Within the dmM, the retrogradely labeled neurons were clustered within the PH next to the 4th ventricle (**Fig. 4c**).

**Figure 4.**
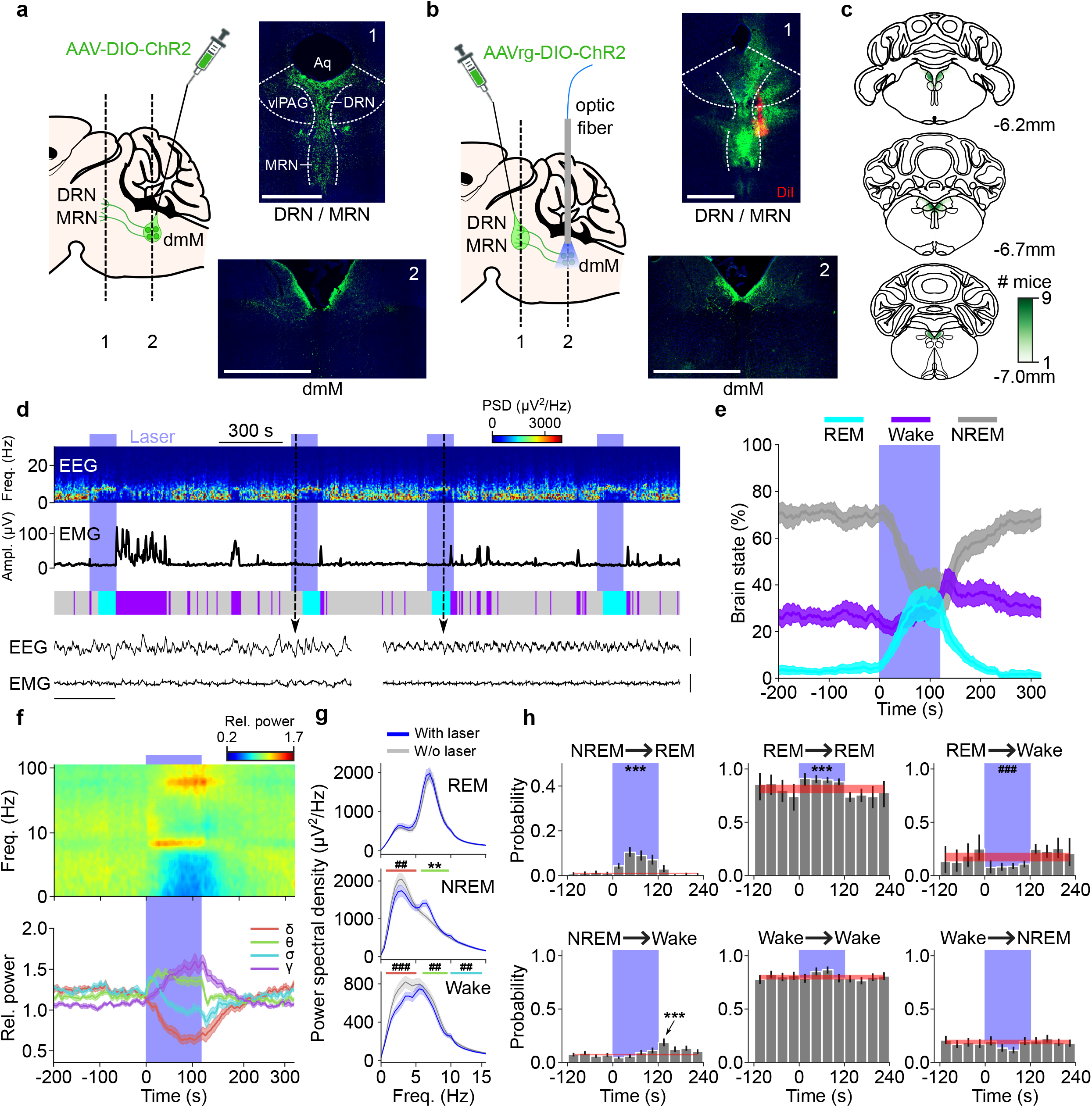
DR/MRN-projecting dmM GAD2 neurons specifically promote REMs. **(a)** Left, schematic sagittal brain section depicting injection of AAV-DIO-ChR2 into the dmM of a GAD2-Cre mouse. Right, fluorescence image of injection site showing expression of ChR2-eYFP in the dmM (bottom) and image showing axonal projections of dmM GAD2 neurons in midbrain and pons (top). Axonal projections were strongest in the dorsal raphe (DRN) and median raphe nucleus (MRN). Blue, Hoechst stain. Scale bars, 1 mm. vlPAG, ventrolateral periaqueductal gray; Aq, aqueduct. **(b)** Schematic depicting injection of AAVrg-DIO-ChR2 into the DR/MRN area of a GAD2-Cre mouse and implantation of an optic fiber into the dmM for stimulation of retrogradely labeled neurons. Top, expression of ChR2-eYFP at the injection area. Bottom, retrogradly labelled neurons in the dmM. Red, DiI labeling solution. Scale bars, 1 mm. **(c)** Expression of ChR2-eYFP in the dmM of GAD2-Cre mice injected with AAVrg-DIO-ChR2 into the DR/MRN. The green color code indicates in how many mice the virus expression overlapped at the corresponding location (n = 9 mice). **(d)** Example experiment. Shown are EEG spectrogram, EMG amplitude, and brain states. Two EEG, EMG raw traces during NREMs and REMs are represented on an expanded timescale for the selected time points (dashed lines; scale bars, 1 s and 0.5 mV). PSD, power spectral density. **(e)** Percentage of REMs, NREMs, and wake before, during, and after laser stimulation (n = 9 mice). Shadings, 95% CIs. Laser stimulation significantly increased REMs (P < 0.001, bootstrap), decreased NREMs (P < 0.001), and did not alter wake (P = 0.32). **(f)** Impact of laser stimulation on EEG spectrogram and different power bands. Top, laser trial averaged EEG spectrogram with logarithmic frequency axis. Each frequency of the spectrogram was normalized by its mean power across the recording. Bottom, delta, theta, sigma, and gamma power before, during, and after laser stimulation. Laser stimulation significantly increased theta (P = 0.007, T = 3.6, paired t-test, n = 9 mice) and gamma power (P = 7.1e-4, T = 5.32), while reducing delta (P = 4.1e-5, T = −8.06) and sigma power (P = 0.004, T = −3.92). Shadings, ± s.e.m. **(g)** Power spectral density of EEG during REMs (top), NREMs (middle), and wake (bottom) with and without laser stimulation. During NREMs optogenetic activation significantly reduced delta (P = 0.0012, T = −5.19; paired t-test, n = 9 mice) and increased theta power (P = 0.0027, T = 4.27). During wake, delta, theta, and sigma power were significantly attenuated (delta, P = 8.3e-4, T = −5.19, theta, P = 0.0018, T = −4.56, sigma, P = 0.0058, T = −3.73; paired t-test). ***/^###^, P < 0.001; **/^##^, P < 0.01 for significant increases/decreases. **(h)** Effect of laser stimulation of DR/MRN-projecting neurons on brain state transition probabilities. Transition probabilities were calculated for 10 s bins. Each bar represents the average transition probability during a 30 s interval. Error bars represent the 95% CI. The red line and shading depict the average baseline transition probability (computed for the interval preceding laser stimulation) and the 95% CI, respectively. Laser stimulation significantly increased the probability of NREMs to REMs transitions (NREM → REM, P < 0.001, bootstrap), REMs to REMs (REM → REM, P < 0.001), while suppressing REMs to wake transitions (REM → Wake, P < 0.001). The probability of wake to wake (Wake → Wake, P = 0.21) and wake to NREMs transitions (Wake → NREM, P = 0.16) was not significantly altered. The arrow indicates a brief increase in NREMs to wake transitions following laser stimulation (P < 0.001). ***/^###^, P < 0.001 for significant increases/decreases.

Optogenetic activation of the DR/MRN-projecting dmM neurons (**Fig. 4d**; 10 Hz, 120 s per trial) strongly increased the percentage of REMs during laser stimulation (**Fig. 4e**; P < 0.001, bootstrap, n = 9 mice) with magnitudes comparable to those observed for stimulation of the whole dmM population (P = 0.16, T = 1.48, unpaired t-test) and also reduced NREMs (P < 0.001). The laser trial averaged EEG spectrogram similarly showed for stimulation of the projection neurons a distinct increase in theta and gamma power with a concomitant reduction in both the delta and sigma power (**Fig. 4f**; delta, P = 4.1e-5, T = −8.06; theta, P = 6.8e-3, T = 3.6; sigma, P = 0.004, T = −3.92; gamma, P = 7.1e-4, T = 5.32, paired t-test). Optogenetic activation during NREMs also suppressed the delta power and increased the theta power (**Fig. 4g**, NREM; delta, P = 0.0012, T = −4.88; theta, P = 0.0027, T = 4.27, paired t-test). Interestingly, in contrast to activation of the whole dmM GAD2 population (**Fig. 1d**), stimulation of the DR/MRN-projecting neurons did not increase the percentage of wakefulness during the laser interval (**Fig. 4e**; P = 0.32, bootstrap) and wake to wake transitions were not significantly altered (**Fig. 4h**; Wake → Wake, P = 0.21, bootstrap). This suggests that the subpopulation of DR/MRN-projecting dmM neurons specifically promotes REMs and does not maintain wakefulness.

Consistent with the strong REMs-promoting effect, activation of the DR/MRN-projecting neurons enhanced both NREMs to REMs and REMs to REMs transitions, demonstrating that these neurons both induce and maintain REMs (**Fig. 4h**; NREM → REM, P < 0.001; REM → REM, P < 0.001). Closed-loop stimulation of the projection neurons during REMs confirmed the REMs-promoting effect (**Suppl. Fig. 3e**; no laser, 55.7 s; laser, 109.8 s; P = 0.022, T = 3.27, paired t-test, n = 6 mice) and REMs periods extended by laser stimulation also increased the total duration of NREMs but not wakefulness during the subsequent inter-REM interval (**Suppl. Fig. 3e**; NREM, P = 0.032, T = 2.95; Wake, P = 0.186, T = 1.53, paired t-test). Taken together, by using an AAV for retrograde labeling of DR/MRN-projecting neurons, we could isolate a subpopulation of dmM GAD2 neurons within the PH that was sufficient for both the initiation and maintenance of REMs.

### dmM GAD2 neurons are strongly activated during REMs and synchronized with sigma power during NREMs

Electrophysiological *in vivo* recordings demonstrated the existence of highly REMs-active neurons in the dmM (Goutagny et al., 2008; Sakai, 2018). Since the dmM contains multiple different cell types (Girard et al., 2020), it is necessary to perform cell-type specific recordings to unravel the state-dependent activity of the GABAergic neurons. To monitor the dynamics of dmM GAD2 neurons *in vivo* during spontaneous sleep, we recorded their population activity using fiber photometry (Lerner et al., 2015). We injected Cre-dependent AAVs encoding the fluorescent calcium indicator GCaMP6s (AAV-Flex-GCaMP6s) into the dmM of GAD2-Cre mice and implanted an optic fiber to measure the calcium dependent fluorescence of GCaMP6s (**Fig. 5a, b; Suppl. Fig. 4b; Suppl. Video 2**). In each mouse, the calcium activity of the dmM GAD2 neurons was significantly modulated by the brain state (**Fig. 5c**; P < 0.05, one-way ANOVA, n = 6 mice). In 5 out of 6 mice, the dmM GAD2 neurons were most active during REMs (P < 0.05, Tukey post-hoc test); in the remaining mouse, the activity was highest during both REMs and wake. Analyzing the calcium changes at brain state transitions, we found that the activity of dmM GAD2 neurons started increasing 35 s before the onset of REMs (**Suppl. Fig. 4a**; P < 0.05, paired t-test with Bonferroni correction). Their activity persisted throughout REMs at high levels and started decreasing 5 s before its termination. This activity profile further suggests that these neurons contribute to REMs initiation and maintenance during natural sleep.

**Figure 5.**
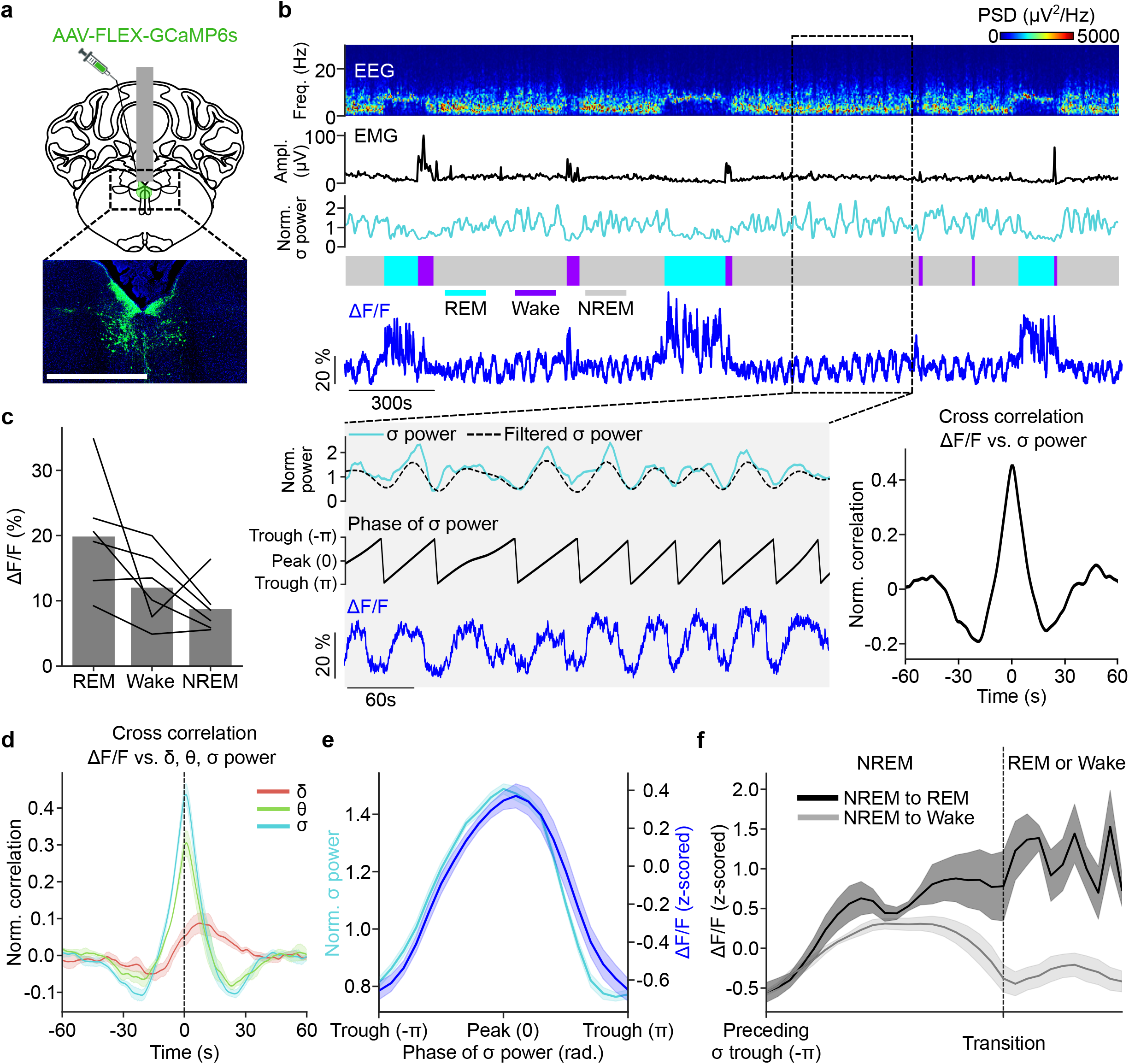
dmM GAD2 neurons are strongly activated during REMs and synchronized with sigma power during NREMs. **(a)** Top, schematic of calcium imaging using fiber photometry. AAVs expressing GCaMP6s were injected into the dmM of GAD2-cre mice. Gray rectangle, optic fiber. Bottom, fluorescence image showing expression of GCaMP6s (green) in the dmM. Blue, Hoechst stain. Scale bar, 1 mm. **(b)** Top, example fiber photometry recording. Shown are EEG spectrogram, EMG amplitude, color-coded brain states, and ΔF/F signal. PSD, power spectral density. Bottom left, the black box indicates an interval for which the EEG sigma power (top, 10 − 15 Hz) and calcium signal (bottom) are shown at an expanded timescale. The filtered sigma power (dashed line) was used to determine the phase of the sigma power oscillation (middle). Bottom right, cross correlation between the calcium signal and sigma power during NREMs for the entire example recording. **(c)** Average ΔF/F activity during REMs, wake, and NREMs. Each line shows the activity of one mouse (n = 6 mice). In each mouse, the brain state significantly modulated the activity (P < 0.05, one-way ANOVA), and in 5 out of 6 animals the calcium activity was highest during REMs (P < 0.05, Tukey post-hoc test). **(d)** Cross correlation between calcium activity and EEG delta, theta, or sigma power during NREMs. The correlation was strongest for the sigma power (P = 1.41e-07, F(2, 10) = 151.47, one-way repeated-measures ANOVA; P < 0.0001, Bonferroni correction).The cross correlation of the two signals was maximal at 0.75 s ± 0.32 s (mean ± s.d.). Shadings, ± s.e.m. The dashed line indicates time point 0 s. **(e)** Average sigma power and calcium activity during a single cycle of the sigma power oscillation. Each sigma power cycle was normalized in time, ranging from −π to π rad. The phase of the sigma power significantly modulated the calcium activity (P = 2.89e-46, F(20, 100) = 58.90, one-way repeated-measures ANOVA). Shadings, ± s.e.m. **(f)** Calcium activity preceding NREMs to REMs or NREMs to wake transitions. The interval from the preceding trough in sigma power until the transition and the duration of the following REMs or wake episode was normalized in time. The activity during NREMs was significantly different depending on whether the mouse transitioned to REMs or wake (F(20,100) = 4.017, P = 0.045, one-way repeated-measures ANOVA; n = 6 mice). Wake also includes microarousals (wake periods ≤ 10s). Shadings, ± s.e.m.

Interestingly, during NREMs the calcium activity of the dmM GAD2 neurons rhythmically fluctuated on an infraslow timescale of tens of seconds (**Fig. 5b**). A recent study showed that the EEG displays a salient oscillation in the sigma power (10 - 15 Hz) on a similar timescale (Lecci et al., 2017). We wondered whether the rhythmic NREMs activity of the dmM inhibitory neurons may follow this infraslow oscillation. We indeed found that the EEG sigma power and calcium activity of dmM GAD2 neurons were strongly positively correlated during NREMs (**Fig. 5b, d; Suppl. Fig. 4c;** P = 1.6e-5, T = 16.17, t-test, n = 6 mice) and the power spectral density of both signals consequently peaked at a comparable frequency (**Suppl. Fig. 4d**, P = 0.695, T = 0.41, paired t-test). The cross correlation of the two signals was maximal at 0.75 s ± 0.32 s (mean ± s.d.) (**Fig. 5d**), indicating that the dmM calcium activity followed the cortical sigma power with short time delay. The correlation between the two signals was strongest during NREMs (**Suppl. Fig. 4c**; P = 2.0e-6, F(2, 10) = 65.32, one-way repeated-measures ANOVA; P < 7.7e-9, Bonferroni correction, n = 6 mice) and the signals were anti-correlated during wakefulness (**Suppl. Fig. 4c**; P = 1.05e-4, T = 16.17, t-test). Compared with the delta and theta power, the correlation was strongest for the sigma power (**Fig. 5c**; one-way repeated-measures ANOVA, P = 1.41e-07, F(2, 10) = 151.47; Bonferroni correction, P < 1.15e-4).

Next, we analyzed the dmM activity throughout single oscillation cycles of the sigma power during consolidated bouts of NREMs (NREMs bouts lasting at least 120 s and only interrupted by microarousals, i.e. wake episodes ≤ 10 s). The ΔF/F activity showed a cosine-shaped activity modulation and closely matched the time course of the sigma power (**Fig. 5e**).

To compare the NREMs activity preceding a wake episode with the activity preceding REMs, we analyzed the calcium signals starting from the last trough in sigma power before the transition to wake or REMs. We found that the activity of dmM GAD2 neurons differed depending on whether the mouse transitioned to wakefulness or REMs (**Fig. 5f**; P = 0.045, F(20,100) = 4.017, n = 6 mice, one-way repeated-measures ANOVA). Preceding a transition to wakefulness, the dmM neuronal population exhibited a cosine-like rise and decay in its activity (**Fig. 5f**). In contrast, before a transition to REMs, the ΔF/F signal raised more steeply and remained elevated before reaching its peak during REMs (**Fig. 5f**), indicating that the activity of the dmM GAD2 population is predictive of whether the animal will transition to REMs or wakefulness.

### Ultradian modulation of dmM GAD2 activity

In addition to the infraslow timescale, we found that the activity of dmM neurons is also modulated on the ultradian timescale. To quantify the ultradian modulation of the calcium signal, we normalized the duration of inter-REM intervals to analyze changes in the dmM activity across these intervals. Consistent with an accumulation of REMs pressure during NREMs, the NREMs calcium activity progressively increased throughout inter-REM intervals (P = 6.85e-7, R = 0.31, linear regression fit), while the wake activity decayed (**Suppl. Fig 4e**; P = 0.006, R = −0.20, linear regression fit). Longer REMs periods are thought to more strongly dissipate REMs pressure. Accordingly, the duration of REMs periods was negatively correlated with the NREMs activity of dmM neurons during the following inter-REM interval (**Suppl. Fig. 4f**; P = 0.029, R = −0.308, linear regression fit); the longer the preceding REMs episode, the less active the dmM neurons during subsequent NREMs. In contrast, the activity during wakefulness was not significantly modulated by the preceding REMs duration (**Suppl. Fig. 4f**; P = 0.846, R = −0.028, linear regression fit), in line with the notion that REMs pressure accumulates during NREMs (Benington and Heller, 1994).

### The sigma power oscillation modulates the delay of optogenetically induced REMs

We showed that the infraslow oscillation in sigma power strongly modulates the activity of dmM neurons. Next, we tested whether this oscillation also plays a role in determining the timing of REMs episodes. In the optogenetic stimulation experiments of dmM GAD2 and DR/MRN-projecting neurons (**Figs. 1, 4**), the delay between laser onset and the start of laser-induced REMs episodes considerably varied across trials (**Fig. 6a**). We wondered whether the phase of the infraslow oscillation at laser onset may influence the latency of laser-induced REMs. As the oscillation in sigma power is particularly pronounced during consolidated NREMs (**Fig. 6b**) (Lecci et al., 2017), we restricted our analysis to laser trials preceded by at least 2 min of NREMs (only interrupted by microarousals) (**Fig. 6c**). We found that the phase of the infraslow oscillation indeed modulated the delay between laser onset and REMs (**Fig. 6d**; P = 0.0017, F(3,48) = 5.85, mixed ANOVA with phase and experiment type (dmM vs. dmM to DR/MRN) as within and between factors), but the delay times did not significantly depend on the experiment type (P = 0.19, F(1,16) = 1.87; **Fig. 6d**). The mean delay time was longest when the onset of laser stimulation coincided with the trough and beginning of the rising phase of the sigma power cycle (-π to −π/2 rad.). In contrast, it was shortest when the laser shortly followed the peak of the infraslow oscillation cycle during the beginning of its falling phase (0 to π/2 rad.), indicating that this phase range establishes an optimal time window during which activation of the dmM GAD2 neurons triggers REMs with minimal latency.

**Figure 6.**
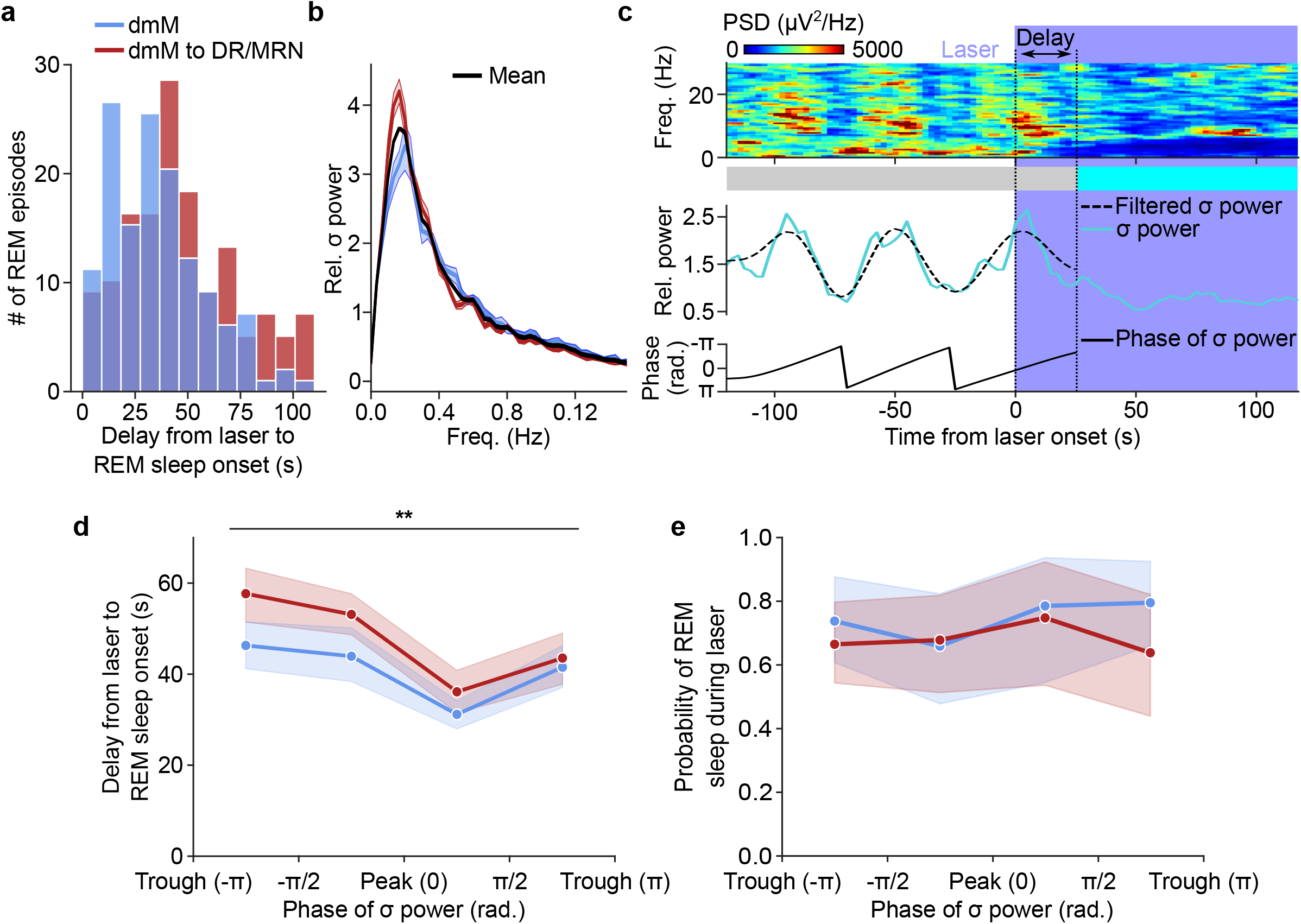
The sigma power oscillation modulates the delay of optogenetically induced REMs. **(a)** Distribution of delay times between onset of laser stimulation and REMs episodes for open-loop stimulation of dmM GAD2 and DR/MRN-projecting neurons during consolidated NREMs (**Figs. 1, 4**; n = 9 mice for each data set). Red, dmM; blue, dmM to DR/MRN. **(b)** Power spectral densities of the EEG sigma power for open-loop stimulation of dmM GAD2 and DR/MRN-projecting neurons during consolidated NREMs. The spectral densities peaked for both data sets in the infraslow range (dmM, 0.021 ± 0.0064 Hz, dmM to DR/MRN, 0.015 ± 0.0029 Hz, mean, 0.018 Hz ± 0.0057; mean ± s.d.), indicating a dominant infraslow oscillation in sigma power with an average wavelength of 55.6 s. For each animal, the spectral density was normalized by its mean power. Shadings, ± s.e.m. **(c)** Example of a laser stimulation trial. Shown are the EEG spectrogram, color coded brain states and sigma power. The filtered sigma power (dashed line) was used to determine the phase of the sigma power oscillation at laser onset. For each laser trial preceded by at least 2 min of NREMs (only interrupted by microarousals), we determined the delay between laser and REMs onset. **(d)** Delay times between laser and REMs onset depending on the phase of the sigma power oscillation at laser onset. Left, delay times for stimulation of dmM GAD2 neurons. Phase values were divided into four equally sized bins. For all laser trials within a given bin, we calculated the average delay time between laser and REMs onset across mice. The delay times were significantly modulated by the phase, but not the experiment type (dmM vs. dmM to DR/MRN) (phase, P = 0.0017, F(3,48) = 5.85; experiment, P = 0.19, F(1,16) = 1.87; interaction - phase * experiment, P = 0.57, F(3, 48) = 0.67; mixed ANOVA with phase and experiment type (dmM vs. dmM to DR/MRN) as within and between factors, n = 9 (dmM) + 9 (dmM to DR/MRN) mice)). **, P < 0.01 for significant modulation by phase. Shadings, ± s.e.m. **(e)** Probability of REMs induction during the 120 s stimulation interval in dependence of the sigma power phase. For each mouse and phase bin, we divided the number of successful trials (where REMs was triggered) by the sum of successful and unsuccessful trials for this bin. In contrast to the delay times, the probability of REMs induction was not significantly modulated (phase, P = 0.64, F(3,48) = 0.64; experiment, P = 0.53, F(1,16) = 0.42; interaction - phase * experiment, P = 0.74, F(3, 48) = 0.42; mixed ANOVA with phase and experiment type as within and between factors). Shadings, 95% CIs.

The finding that the delay time depends on the phase suggests that the oscillation in sigma power indeed modulates the timing of the induction of REMs. In contrast to the delay, the probability of REMs induction was not significantly modulated by the sigma power phase (**Fig. 6e**; phase, P = 0.64, F(3,48) = 0.57; experiment type, P = 0.53, F(1,16) = 0.42, mixed ANOVA with phase and experiment type as within and between factors), likely because the 2 min laser stimulation interval offered sufficient opportunity for the laser to eventually overlap with the optimal sigma power range. Hence, our results suggest that while optogenetic activation of dmM neurons strongly increases the probability that REMs occurs during the 2 min laser stimulation interval (**Fig. 1d, Fig. 4e**), the infraslow oscillation contributes to determining when the REMs episode is initiated (**Fig. 6d**).

## DISCUSSION

Using optogenetic manipulation, we found that GAD2 neurons in the dmM promote the initiation of REMs and maintain both REMs and wakefulness (**Figs. 1, 2**), while chemogenetic inhibition suppresses REMs (**Fig. 3**). Viral anterograde tracing showed that the dmM neurons strongly project to the DRN and MRN. Using an AAV variant with high retrograde efficiency, we specifically expressed ChR2-eYFP in the subpopulation of dmM neurons projecting to the DR/MRN (**Fig. 4**), which were mainly located within the PH next to the 4th ventricle. Optogenetic activation of these projection neurons enhanced both the initiation and maintenance of REMs without maintaining wakefulness, suggesting that they specifically promote REMs. Using fiber photometry, we showed that dmM GAD2 neurons are highly active during REMs, consistent with their role in promoting REMs (**Fig. 5**). Interestingly, during NREMs the activity of these neurons slowly fluctuates in close synchrony with the EEG sigma power. The infraslow oscillation in sigma power influenced the latency with which optogenetic activation of dmM neurons triggered REMs (**Fig. 6**), suggesting that it plays an important role in timing REMs episodes.

### GABAergic medulla neurons promote REMs through inhibition of REMs-off neurons in pons and midbrain

Similar to the dmM, GAD2 neurons in the ventral medulla (vM) have also been shown to strongly promote REMs (Weber et al., 2015). The vM REMs-promoting neurons innervate vlPAG GABAergic neurons (Weber et al., 2015) and in contrast to the dmM neurons project less densely to the raphe nuclei (Gervasoni et al., 2000). Comparable to the effects observed for chemo- or optogenetic excitation of GABAergic neurons in the vlPAG (Hayashi et al., 2015; Weber et al., 2018), a recent study showed that tonic optogenetic activation of serotonergic neurons in the DRN also suppresses REMs, while maintaining NREMs (Oikonomou et al., 2019). Fiber photometry imaging, consistent with previous electrophysiological recordings (McGinty and Harper, 1976), demonstrated that their activity is lowest during REMs. Hence, inhibitory neurons in the dmM and vM may constitute two complementary circuit nodes that act in concert to initiate and maintain REMs by suppressing REMs-off neurons throughout the midbrain and pons, including neurons in the DRN, MRN, and vlPAG. In addition to these areas, the LC also receives strong axonal projections from inhibitory neurons in PH and DPGi (Ennis and Aston-Jones, 1989; Verret et al., 2006). Pharmacological inhibition of the LC suppressed the extension of REMs episodes elicited by electrical stimulation of the PH, suggesting that this projection contributes to REMs maintenance (Kaur et al., 1997, 2001). As previous studies also demonstrated the importance of the DPGi in regulating REMs (Clément et al., 2014; Goutagny et al., 2008; Verret et al., 2006), it would be interesting to similarly test the specific role of DPGi inhibitory neurons using projection targets preferentially innervated by these neurons. Furthermore, identification of specific cell markers differentiating PH from DPGi neurons would allow for more targeted approaches in delineating the connectivity and function of these adjacent medullary nuclei in REMs control (Girard et al., 2020; Gutierrez Herrera et al., 2019).

Stimulation of dmM GAD2 neurons strongly promoted theta oscillations during NREMs indicating that these neurons directly or indirectly interact with circuits involved in generating hippocampal theta oscillations. Pharmacological inhibition (Kinney et al., 1994, 1995; Vertes et al., 1994) and lesions of the MRN (Maru et al., 1979) elicit theta oscillations. This effect is likely mediated by serotonergic neurons, as pharmacological stimulation of serotonergic autoreceptors in the MRN, which inactivates serotonergic neurons, triggers theta oscillations (Vertes et al., 1994). Postsynaptic inhibition of the MRN by the dmM may therefore contribute to the induction of theta oscillations before and during REMs and underlie the increased EEG theta power during NREMs when stimulating dmM neurons.

In addition to circuits directly regulating REMs, the medulla contains further neural populations that are involved in generating defining properties of REMs. Glycinergic neurons in the ventromedial medulla have been shown to be crucial for the characteristic paralysis of skeletal muscles during REMs (Chase et al., 1989; Holstege and Bongers, 1991; Schenkel and Siegel, 1989; Valencia Garcia et al., 2018; Uchida et al., 2020) and calbindin-expressing neurons within the DPGi are necessary for the occurence of eye movements during REMs (Gutierrez Herrera et al., 2019). The medulla thus harbors a hub of circuits for the control of REMs and its defining electrophysiological and behavioral features.

### Neural correlates of the infraslow oscillation

Using fiber photometry imaging, we found that the activity of dmM neurons is strongly modulated by the infraslow oscillation in sigma power. A previous study showed that this infraslow oscillation modulates the arousability of mice during NREMs. When presented with acoustic stimuli during NREMs, mice tended to sleep through when the stimulus coincided with the rising phase, but to wake up during the falling phase (Lecci et al., 2017). We found that the phase of the infraslow oscillation influenced the delay of optogenetically induced REMs episodes, suggesting that it also plays an important role in the induction of REMs. The infraslow oscillation may directly contribute to the timing of NREMs to REMs transitions by modulating the activity of REMs regulatory neurons, as we have observed for the dmM neurons, providing a link between spontaneous brain activity fluctuations on the infraslow timescale and brain state transitions. While our analysis demonstrates that the falling phase offers an optimal time window to trigger REMs with minimal latency during the 2 min long laser stimulation interval, more temporally precise manipulation of the dmM neurons could allow for testing whether the infraslow oscillation also determines the probability of REMs induction, as has been shown for awakenings from NREMs (Fernandez and Lüthi, 2019; Lecci et al., 2017).

The neural correlates underlying the infraslow oscillation are still largely unknown. This oscillation is present in the EEG / local field potential across the whole cortical surface (Lecci et al., 2017). Consistent with the fact that sleep spindles are a major contributor to the sigma power (Fernandez et al., 2018), multi-unit activity in the thalamus, which is critical for spindle generation, is modulated on the infraslow timescale (Csernai et al., 2019). The calcium activity in apical dendrites of cortical layer 5 neurons is also tightly correlated with sigma power oscillations (Seibt et al., 2017). Together with our study, these findings suggest that this oscillation is a global rhythm shaping the spontaneous activity across multiple brain areas. As the calcium activity of the dmM GAD2 neurons lags behind the cortical sigma power (**Fig. 5d, e**), it is feasible that dmM neurons receive direct or indirect inputs from the thalamocortical system, possibly driving their oscillatory activity during NREMs. Such inputs could also explain why enhancing the spindle rate through optogenetic activation of the thalamic reticular nucleus increases transitions to REMs (Bandarabadi et al., 2020).

Besides neural activity, the intrinsic blood oxygen level dependent (BOLD) signals in humans and the body temperature also exhibit modulations on the infraslow timescale (Csernai et al., 2019; Mitra et al., 2015) and autonomic variables such as heart rate and pupil diameter are closely correlated with the sigma power oscillation (Lecci et al., 2017; Yüzgeç et al., 2018). It would therefore be interesting for future studies to test whether the dmM neurons interact with output circuits of the autonomic nervous system to coordinate infraslow modulations in neural activity with fluctuations in autonomic activity, which are characteristic of mammalian sleep.

## METHODS

### Animals

All experimental procedures were approved by the Institutional Animal Care and Use Committee (IACUC) at the University of Pennsylvania and conducted in accordance with the National Institutes of Health Office of Laboratory Animal Welfare Policy. Experiments were performed in male or female GAD2-IRES-Cre mice (Jackson Laboratory stock no. 010802). Animals were housed on a 12-h dark/12-h light cycle (lights on between 7 am and 7 pm) and were aged 6-12 weeks at the time of surgery. Male and female mice were distributed equally across groups in each experiment. All mice were group-housed with ad libitum access to food and water.

### Surgical Procedures

All surgeries were performed following the IACUC guidelines for rodent survival surgery. Prior to surgery, mice were given meloxicam subcutaneously (5 mg/kg). Mice were anesthetized using isoflurane (1 - 4%) and positioned in a stereotaxic frame. Animals were placed on a heating pad to maintain the body temperature throughout the procedure. Following asepsis, the skin was incised to gain access to the skull. For virus injection or implantation, a small burr hole was drilled above the dorsomedial medulla (anteroposterior (AP) −6.4 to −6.7 mm, mediolateral (ML) 0 mm).

For optogenetic activation of dmM GAD2 neurons, 0.1 to 0.3 μl of AAV1-EF1a-DIO-hChR2-eYFP-WPRE-hGH (University of Pennsylvania vector core) was injected into the target area of GAD2-Cre mice using Nanoject II (Drummond Scientific) via a glass micropipette (dorsoventral (DV) −3.6 mm). For controls, we injected 0.1 to 0.3 μl of AAV2-Ef1a-DIO-eYFP (University of Pennsylvania vector core) into the same area. To optogenetically stimulate dmM GAD2 neurons projecting to the DR/MRN, an additional burr hole was drilled (AP −4.8 mm, ML 0.0 to −0.25 mm) and 0.1 to 0.25 μl of AAVrg-EF1a-DIO-hChR2-eYFP-WPRE-HGHpA (Addgene) were injected into the DR/MRN (DV −3.2 to −3.4 mm). To localize the injection site, the micropipette was coated with red DiI (Invitrogen Vybrant DiI Cell Labeling Solution, V22885). After virus injection, an optical fiber (0.2 mm diameter) was inserted into the dmM (DV −3.4 to −3.5 mm). For fiber photometry experiments 0.1 to 0.3 μl of AAV1-Syn-Flex-GCaMP6s-WPRE-SV40 (University of Pennsylvania vector core) was injected into dmM and the optic fiber (0.4 mm diameter) was placed on top of the injection site (DV −3.3 to −3.5 mm). For chemogenetic inhibition experiments, 100-200 nl of AAV8-hSyn-DIO-hM4D(Gi)-mCherry (Addgene) were injected (DV −3.6 mm, ML 0 mm). Control mice were instead injected with AAV8-hSyn-DIO-mCherry (Addgene).

EEG signals were recorded using stainless steel wires attached to two screws, one on top of the hippocampus and one on top of the prefrontal cortex. The reference screw was inserted on top of the left cerebellum. For EMG recordings, two stainless steel wires were inserted into the neck muscles. All electrodes, screws, connectors, and optic fibers were secured to the skull using dental cement. After injection and implantation were finished, bupivacaine (2 mg/kg) was administered at the incision site. For fiber photometry, opto-, and chemogenetic experiments, we excluded animals where no virus expression could be detected or where the virus expression extended below the dmM, as were mice in which the optic fiber tip was located below the virus expression.

### Histology

Mice were deeply anesthetized and transcardially perfused with 0.1 M phosphate-buffered saline (PBS) followed by 4% paraformaldehyde in PBS. After removal, brains remained overnight in fixative and were then stored in 30% sucrose by volume in PBS solution for at least one night. After embedding and freezing, brains were sliced into 30 or 40 μm sections using a cryostat (Thermo Scientific HM525 NX) and mounted onto glass slides.

For immunohistochemistry, brain sections were washed in PBS, permeabilized using PBST (0.3% Triton X-100 in PBS) for 30 minutes and then incubated in blocking solution (5% normal donkey serum in PBST) for 1 hour. To stain eYFP-expressing axon fibers, brain sections were subsequently incubated with a chicken anti-GFP primary antibody (Aves Lab, GFP8794984, 1:1,000) diluted in PBS for one night at 4°C. The next day, brain sections were incubated for 2 hours with a species-specific secondary antibody conjugated with green Alexa fluorophore (Jackson ImmunoResearch Laboratories, Inc., 703-545-155, 1:500; donkey anti-chicken) diluted in PBS. The slices were washed with PBS followed by counterstaining with Hoechst solution (33342, Thermo Scientific) and coverslipped with Fluoromount-G (Southern Biotechnic). Fluorescence images were taken using a fluorescence microscope (Microscope, Leica DM6B; Camera, Leica DFC7000GT; LED, Leica CTR6 LED).

### Fitting Histology to Reference Images

To generate heatmaps of virus expression across mice (**Figs. 1b, 3b, 4c**), coronal reference images were downloaded from Allen Mouse Brain Atlas for the appropriate AP coordinates (© 2015 Allen Institute for Brain Science. Allen Brain Atlas API. Available from: http://brain-map.org/api/index.html). For a given AP reference section, the corresponding histology section from each mouse was overlaid and fitted to the reference. Regions in which eYFP-labeled cell bodies could be detected were then outlined by hand. Custom Python programs then detected these outlines and determined for each location on the reference picture the number of mice with overlapping virus expression, which was encoded using different green color intensities.

### Polysomnographic Recordings

Sleep recordings were performed in the animal’s home cage or in a cage to which the mouse had been habituated for 3 days, which was placed within a sound-attenuating box. For opto- and chemogenetic studies, EEG and EMG signals were recorded using an RHD2000 amplifier (intan, sampling rate 1 kHz). For fiber photometry, we used a TDT RZ5P amplifier (sampling rate 1.5 kHz). EEG and EMG signals were referenced to a common ground screw, placed on top of the cerebellum. Videos were recorded using a camera (FLIR, Chameleon3 or ELB, Mini USB camera) placed above the mouse cage. During the recordings, EEG and EMG electrodes were connected to flexible recording cables using a small connector. To determine the brain state of the animal, we first computed the EEG and EMG spectrogram with consecutive fast fourier transforms (FFTs) calculated for sliding, half-overlapping 5 s windows, resulting in 2.5 s time resolution. Next, we computed the time-dependent delta, theta, sigma, and high gamma power by integrating frequencies in the range 0.5 to 4 Hz, 5 to 12 Hz, 12 to 20 Hz, and 100 to 150 Hz, respectively. We also calculated the ratio of the theta and delta power (theta/delta) and the EMG power in the range 50 to 500 Hz. For each power band, we used its temporal mean to separate it into a low and high part (except for the EMG and theta/delta ratio, where we used the mean plus one standard deviation as threshold). REMs was defined by high theta/delta ratio, low EMG, and low delta power. A state was set as NREMs, if delta power was high, the theta/delta ratio was low and EMG power was low. In addition, states with low EMG power, low delta, but high sigma power were scored as NREMs. Wake encompassed states with low delta power and high EMG power and each state with high gamma power (if not otherwise classified as REMs). Finally, we manually verified the automatic classification using a graphical user interface, visualizing the raw EEG, EMG signals, EEG spectrogram, EMG amplitude, and hypnogram. The software for automatic brain state classification and manual inspection was programmed in Python.

### Optogenetic Manipulation

We performed optogenetic experiments 3 to 6 weeks after surgery. Animals were habituated for at least two days to the recording setup. After habituation, sleep recordings for optogenetics were performed during the light cycle (between 8 am and 6 pm) and lasted 6.5 hours on average. For optogenetic experiments, mice were tethered to an optic fiber patch cable in addition to the cable used for EEG/EMG recordings. For optogenetic open-loop stimulation, we repeatedly presented 10 Hz pulse trains (10 ms up, 90 ms down) lasting for 120 s generated by a blue 473 nm laser (4 - 6 mW, including closed-loop stimulation; Laserglow). The inter-stimulation interval was randomly chosen from a uniform distribution ranging from 10 to 20 min. TTL pulses to trigger the laser were controlled using a raspberry pi, which in turn was controlled by a custom-programmed user interface programmed in Python. For optogenetic closed-loop stimulation, the program determined whether the animal was in REMs or not based on real-time spectral analysis of the EEG and EMG signals. The onset of REMs was defined as the time point where the EEG theta/delta ratio exceeded a hard threshold (mean + std of theta/delta), which was calculated using previous recordings from the same animal. REMs lasted until the theta/delta ratio dropped below a soft threshold (mean of theta/delta) or if the EMG amplitude passed an offline calculated threshold. When REMs was detected, the laser turned on with 50% probability and turned off when the REMs episode ended. Optogenetic sleep recordings which contained strong EEG or EMG artifacts were excluded. For EEG power spectral density analysis (**Fig. 3g**) and EMG amplitude analysis (**Suppl. Fig. 3d**) we excluded one recording due to artifacts in the EEG or EMG signal, respectively. These recordings were still included in the remaining analyses, because the artifacts did not affect sleep state scoring.

### Chemogenetic Manipulation

On recording days, saline or CNO (5 mg/kg, Clozapine N-oxide dihydrochloride, Tocris, Cat. No. 6329) was injected intraperitoneally (i.p.) into GAD2-Cre mice expressing hM4D(Gi) or mCherry in the dmM. Each recording session started 20 min after injection and each animal contributed 2 saline and 2 CNO recordings sessions to the data set. For statistical analysis, we compared the impact of CNO or saline in hm4Di or mCherry-expressing mice on the first 4 hours of sleep using mixed ANOVA with virus (mCherry vs. hM4D(Gi)) and intervention (saline vs CNO) as within and between factors, followed by pairwise t-tests with Bonferroni correction.

### Fiber Photometry

Calcium imaging using fiber photometry was performed in mice freely moving in their home cages, placed within a sound-attenuating chamber. The implanted optic fiber was connected to a flexible patch cable. In addition, a flexible cable was connected to the EEG/EMG electrodes via a mini-connector. For calcium imaging, a first LED (Doric Lenses) generated the excitation wavelength of 465 nm and a second LED emitted 405 nm light, which served as control for bleaching and motion artifacts, as the emission signal from the 405 nm illumination is independent of the intracellular calcium concentration. The 465 and 405 nm signals were modulated at two different frequencies (210 and 330 Hz). Both lights were passed through dichroic mirrors before entering a patch cable attached to the optic fiber. Fluorescence signals emitted by GCaMP6s were collected by the optic fiber and passed via the patch cable through a dichroic mirror and GFP emission filter (Doric Lenses) before entering a photoreceiver (Newport Co.). Photoreceiver signals were relayed to an RZ5P amplifier (TDT) and demodulated into two signals using TDT’s Synapse software, corresponding to the 465 and 405 nm excitation wavelengths. For further analysis, we used custom-written Python scripts. First, both signals were low-pass filtered at 2 Hz using a 4th order digital Butterworth filter. Next, using linear regression, we fitted the 405 nm to the 465 nm signal. Finally, the linear fit was subtracted from the 465 nm signal (to correct for photo-bleaching or motion artifacts) and the difference was divided by the linear fit yielding the ΔF/F signal. To determine the brain state, EEG and EMG signals were recorded together with fluorescence signals using the RZ5P amplifier. All recordings were performed during the light phase between 8am and 4pm and lasted for 2 hours. We excluded fiber photometry recordings which contained sudden shifts in the baseline (likely due to a loose connection between optic fiber and patch cord), or with only one or no REMs episodes (as we could not perform inter-REM interval analyses with these recordings).

### Analysis of Infraslow Sigma Power Oscillations

To calculate the power spectral density of the EEG sigma power (**Fig. 6b, Suppl. Fig. 4d**), we first calculated for each recording the EEG power spectrogram by computing the FFT for consecutive sliding, half-overlapping 5 s windows. Next, we normalized the spectrogram by dividing each frequency component by its mean power and calculated the normalized sigma power by averaging across the spectral density values in the sigma range (10 to 15 Hz). As the infraslow rhythm is most pronounced in consolidated NREMs bouts (Lecci et al., 2017), we only considered NREMs bouts that lasted at least 120 s and were only interrupted by microarousals (wake periods ≤ 10s). We then calculated the power spectral density using Welch’s method (with Hann window) for each consolidated NREMs bout and averaged for each animal across the resulting densities.

To determine the instantaneous phase of the infraslow rhythm, we smoothed the sigma power using a 10 s box filter and band-pass filtered it in the range of 0.01 - 0.03 Hz using a 4th order digital Butterworth filter. Finally, we computed the phase angle by applying the Hilbert transform to the band-pass filtered sigma power signal (**Figs. 5b, 6c**). Based on the phase, we could then isolate the beginning and end of single infraslow cycles to average the ΔF/F activity of dmM neurons during single cycles (**Fig. 5e**) or to determine the phase at the onset of laser stimulation in the optogenetic open-loop experiments (**Fig. 6**). To calculate the ΔF/F activity during single cycles, we first downsampled the calcium signals to the same temporal resolution (2.5 s) as the sigma power, and then normalized the cycle durations for averaging.

To compute Pearson’s r between the ΔF/F signals and the normalized sigma power (or other power bands) (**Suppl. Fig. 4c**), we correlated for each mouse all states scored as REMs, NREMs, or wake with the corresponding 2.5 s bins in the calcium signal.

To calculate the cross correlation between ΔF/F signals and sigma power (or other power bands) (**Fig. 5b, d**), we first calculated for all NREMs bouts equal or larger than 120 s (possibly interrupted by microarousals) the sigma power (**s**) from the EEG spectrogram, computed using consecutive 2.5 s windows, with 80% overlap to increase the temporal resolution. We again normalized the spectrogram by dividing each frequency component by its mean power. Using the same overlapping binning, we downsampled the ΔF/F signal (**d**) to prevent any time lags resulting from differences in downsampling and then calculated the cross correlation of both signals. The cross correlation was normalized by dividing it by the product of the standard deviation of **s** and **d** and the number of data points in **s** and **d**. For each mouse we finally obtained the mean cross correlation by averaging across all NREMs bouts.

### Statistics

Statistical analyses were performed using the python modules scipy.stats (scipy.org) and pingouin (https://pingouin-stats.org). We did not predetermine sample sizes, but cohorts were similarly sized as in other relevant sleep studies (Eban-Rothschild et al., 2016; Yu et al., 2019). All statistical tests were two-sided. The significance of changes in brain state percentages or transition probabilities between brain states induced by laser stimulation were tested using bootstrapping. Otherwise, data were compared using one-way or two-way ANOVA followed by multiple comparisons tests or t-tests. We verified that the data were normally distributed using the Shapiro-Wilk test for normality. For percentages and probabilities, we computed 95% confidence intervals using bootstrapping and visualized them using custom-written functions or seaborn (https://seaborn.pydata.org). Otherwise, results in figures were represented as mean ± s.e.m.

## Acknowledgements

This work was supported by the National Institute of Health (NIH)/National Heart, Lung, and Blood Institute (NHLBI), R01HL149133 to FW, a NARSAD Young Investigator grant (#27799) to FW by the Brain & Behavior Research Foundation, and by a grant from the Margaret Q. Landenberger Foundation to FW. We thank J. Hong for help with sleep recordings and J. Smith and X. Li for help with histology.

## Author Contributions

JAS, SC, and FW conceived and designed the study. JAS performed all optogenetic and fiber photometry experiments and analyzed all sleep data. ALS performed optogenetic pilot experiments. JB built the setup for optogenetic sleep recordings including the software to run experiments and the setup for fiber photometry experiments. JAS and FW analyzed the data and wrote the manuscript.

---

**Supplementary Figure 1.**
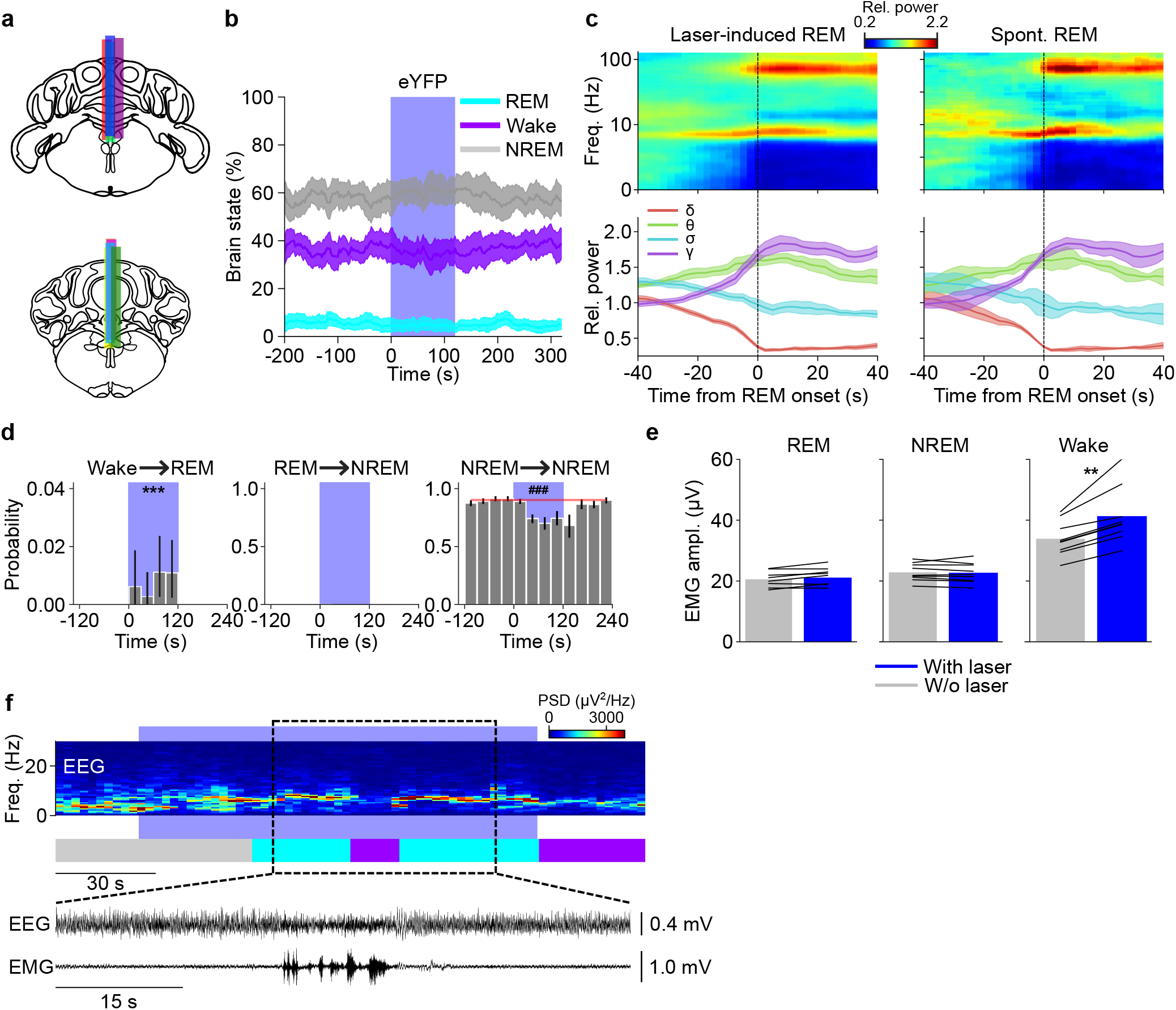
Effects of laser stimulation of dmM GAD2 neurons on EEG and EMG in mice expressing ChR2-eYFP or eYFP. **(a)** Location of fiber tracts for optogenetic stimulation experiments. Each colored bar represents the location of an optic fiber used for optogenetic stimulation of dmM GAD2 neurons (n = 9 mice). The coronal brain schemes were adapted from Allen Mouse Brain Atlas (©2015 Allen Institute for Brain Science. Allen Brain Atlas API. Available from: http://brain-map.org/api/index.html). **(b)** Effect of laser stimulation on REMs, NREMs, and wake in eYFP control mice (n = 6). Laser stimulation did not significantly change the percentage of any brain state (P > 0.233, bootstrap). **(c)** Comparison of the EEG spectrogram for laser-induced and spontaneous NREMs to REMs transitions. Left, mean EEG spectrogram before and after a NREMs to REMs transition, averaged across all laser-induced REMs episodes. Each frequency component of the spectrogram was normalized by its mean power across the recording. The mean time course of delta (0.5 - 4.5 Hz), theta (6 - 9 Hz), sigma (10 −15 Hz), and gamma (55 − 90 Hz) power is shown at the bottom. Right, mean EEG spectrogram for spontaneous REMs episodes (not overlapping with laser stimulation) along with the mean time course of delta, theta, sigma, and gamma power during the transition. Time point 0 s corresponds to the transition point. Shadings, *±* s.e.m. **(d)** Effect of laser stimulation on wake to REMs, REMs to NREMs, and NREMs to NREMs transition probabilities. In rare instances, laser stimulation triggered wake to REMs transitions (Wake → REM, P < 0.001, bootstrap; see (**f**) for an example). We did not observe any REMs to NREMs transitions during the baseline and laser stimulation interval. The strong increase in NREMs to REMs transitions by optogenetic activation (**Fig. 1g**) resulted in a reduction of NREMs to NREMs transitions (NREM → NREM, P < 0.001), i.e. the maintenance of NREMs was impaired during laser stimulation. The red line and shading depict the average baseline transition probability (computed for the interval preceding laser stimulation) and the 95% CI, respectively. Bars, average transition probabilities; error bars, 95% CIs. ^∗∗∗^*/*^###^, P < 0001 for significant increases/decreases. **(e)** EMG amplitude during REMs, NREMs, and wake with and without (w/o) laser stimulation. Optogenetic activation of dmM GAD2 neurons increased the EMG amplitude during wakefulness, but did not affect the amplitude during REMs, or NREMs (REM, P = 0.108, T = 1.84; NREM, P = 0.751, T = −0.33; Wake, P = 3.7e-3, T = 4.05, paired t-test). The left plot (REM) shows data for only 8 mice, as one mouse had no spontaneous REMs episodes. Bars, average across mice; lines, individual mice; ^∗∗^, P < 0.01. **(f)** Example of a wake to REMs transition. Shown are EEG spectrogram and hypnogram. The black box indicates an interval for which the EEG and EMG are shown at an expanded timescale. For each observed wake to REMs transition, the wake episode was directly preceded by REMs. In total, we observed 16 laser-induced wake to REMs transitions in 6 out of n = 9 mice. PSD, power spectral density.

**Supplementary Figure 2.**
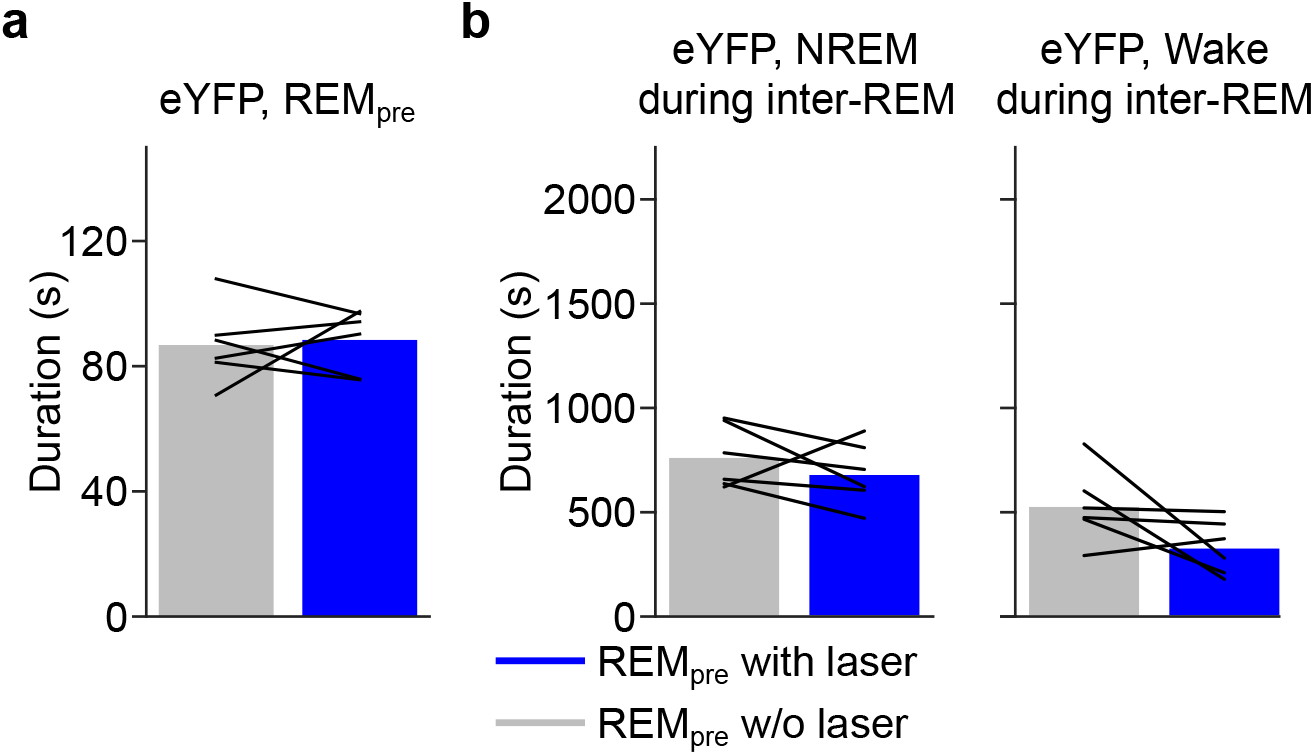
Closed-loop stimulation in eYFP control mice. **(a)** Closed-loop laser stimulation in eYFP control mice did not significantly alter the duration of REMs episodes (P = 0.80, paired t-test, n = 6 mice). Bars, average across mice; lines, individual mice. **(b)** Effects of laser stimulation on NREMs and wake during inter-REM. Left, the total duration of NREMs during inter-REM did not depend on whether the preceding REMs episode overlapped with laser stimulation or not (P = 0.351, paired t-test). Right, total duration of wake during inter-REM (P = 0.109, paired t-test). Bars, average across mice; lines, individual mice.

**Supplementary Figure 3.**
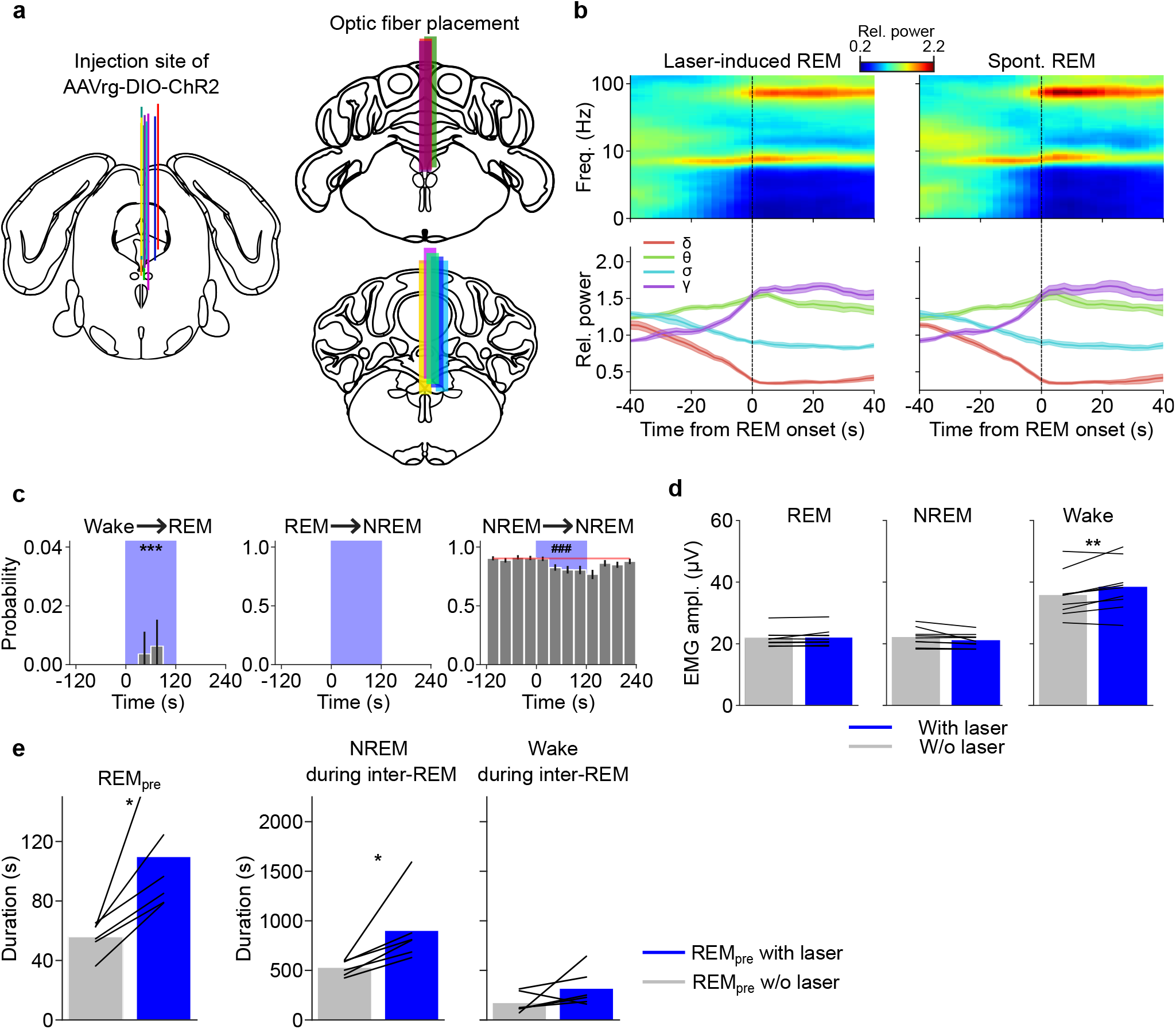
Effects of open and closed-loop stimulation of DR/MRN-projecting dmM GAD2 neurons. **(a)** Location of injection sites of AAVrg-DIO-ChR2 into the DR/MRN (left) and placement of optic fibers for optogenetic stimulation in the dmM (right). The location of the injection site was identified using DiI. **(b)** Comparison of the EEG spectrogram for laser-induced and spontaneous NREMs to REMs transitions. Left, mean EEG spectrogram before and after a NREMs to REMs transition, averaged across all laser-induced REMs episodes. Each frequency component of the spectrogram was normalized by its mean power across the recording. The mean time course of different power bands (delta, theta, sigma, and gamma) is shown at the bottom. Right, mean EEG spectrogram for spontaneous REMs episodes (not overlapping with laser stimulation) along with the mean time course of different power bands during the transition. Time point 0 s corresponds to the transition point. Shadings, *±* s.e.m., n = 9 mice. **(c)** Effect of laser stimulation on wake to REMs, REMs to NREMs, and NREMs to NREMs transition probabilities. Similar to stimulating the whole population of dmM GAD2 neurons (**Suppl. Fig 1d**,**f**), laser stimulation of the DR/MRN-projecting neurons triggered wake to REMs transitions in rare instances (Wake → REM, P < 0.001, bootstrap, n = 9 mice). In total, we observed 4 wake to REMs transitions in 3 out of n = 9 mice. We did not observe any REMs to NREMs transitions during baseline or laser stimulation intervals. Similar to stimulation of the whole population of dmM GAD2 neurons (**Suppl. Fig 1d**), activation of the DR/MRN-projecting neurons caused a reduction of NREMs to NREMs transitions (NREM → NREM, P < 0.001), reflecting an impaired maintenance of NREMs. The red line and shading depict the average baseline transition probability (computed for the interval preceding laser stimulation) and the 95% CI, respectively. Bars, average transition probabilities; error bars, 95% CIs. ^∗∗∗^*/*^###^, P < 0001 for significant increases/decreases. **(d)** EMG amplitude during REMs, NREMs, and wake with and without laser stimulation. Optogenetic activation of DR/MRN-projecting neurons increased the EMG amplitude during wakefulness, but did not affect the amplitude during REMs, or NREMs (REM, P = 0.238, T = 1.27; NREM, P = 0.115, T = −1.77; Wake, P = 0.005, T = 3.80, paired t-test). ^∗∗^, P < 0.01. Bars, average across mice; lines, individual mice. **(e)** Results for closed-loop stimulation of DR/MRN-projecting dmM GAD2 neurons. Left, duration of REMs episodes with and without laser. Closed-loop stimulation significantly prolonged the duration of REMs episodes (P = 0.022, T = 3.27, paired t-test, n = 6 mice). ^∗^, P < 0.05. REMs episodes overlapping with laser stimulation were followed by a larger total duration of NREMs during the subsequent inter-REM interval (P = 0.032, T = 2.95, paired t-test), while the total duration of wakefulness was not significantly altered (P = 0.186, T = 1.52, paired t-test). Bars, average across mice; lines, individual mice. ^∗^, P < 0.05.

**Supplementary Figure 4.**
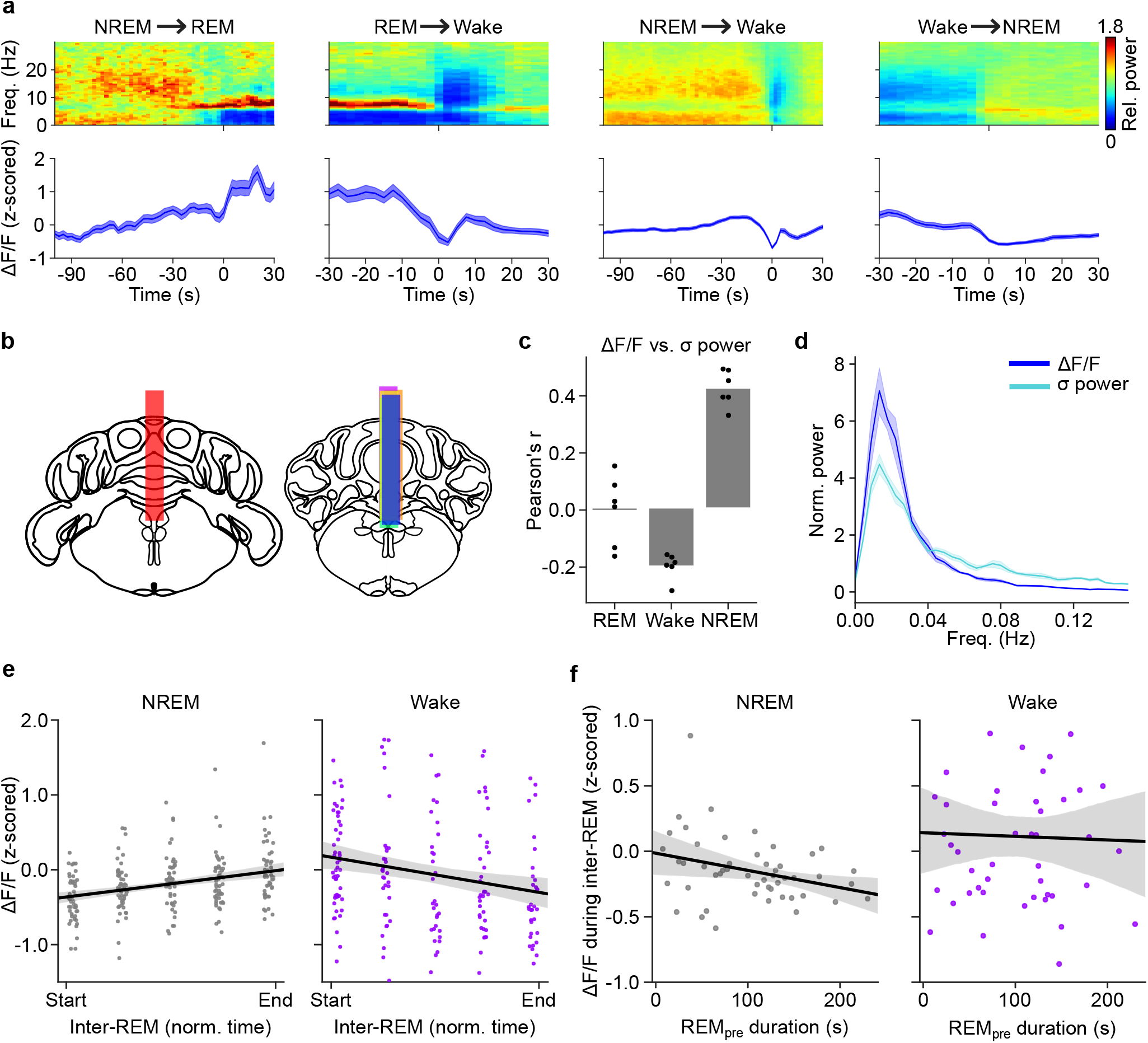
Calcium activity of dmM GAD2 neurons during state transitions, NREMs, and inter-REM. **(a)** Calcium activity (ΔF*/*F, z-scored) at brain state transitions. For each transition from state X to Y, we ensured that the mouse was in state X for at least 100 s (NREM → REM and NREM → Wake) or 30 s (REM → Wake and Wake → NREM). For NREMs to REMs transitions, the activity was significantly increased 35 s before the actual transition (P < 0.05, paired t-test with Bonferroni correction for n = 62 NREMs to REMs transitions from n = 6 mice). We compared the activity in the baseline interval ranging from −100 s to −90 s to consecutive 10 s bins. For REMs to wake transitions, the calcium activity significantly dropped 5 s before the transition (n = 63 transitions). For NREMs to wake transitions, the activity started significantly increasing 35 s before the transition (n = 252 transitions). For wake to NREMs transitions, the activity was significantly reduced 15 s before the transition (n = 186 transitions). Shadings, *±* s.e.m. **(b)** Each colored bar represents the location of an optic fiber implant for photometry imaging. **(c)** Pearson correlation between ΔF*/*F and sigma power for REMs, wake, and NREMs. The correlation between the two signals was highest during NREMs (P = 1e-6, F(2,10) = 70.73, one-way repeated measures ANOVA; P < 1e-5, Bonferroni correction, n = 6 mice). Dots, individual mice. **(d)** Normalized spectral density of sigma power and ΔF*/*F during NREMs. The power spectral density for both signals was calculated for consolidated bouts of NREMs (episodes ≥ 120 s, only interrupted by microarousals, i.e. wake episodes ≤ 10 s) and normalized for each animal by its mean power. The peak frequencies for ΔF*/*F and sigma power were 0.0156 Hz *±* 0.0037 Hz (mean *±* s.d.) and 0.0163 Hz *±* 0.0036 Hz and were not significantly different (P = 0.36, T = 0.42, paired t-test, n = 6 mice). Shadings, *±* s.e.m. **(e)** Calcium activity (ΔF*/*F, z-scored) during NREMs and wake states within different segments of the inter-REM interval. Each inter-REM interval (interval between two successive REMs episodes) was normalized in time and divided into five consecutive bins. The activity during NREMs steadily increased throughout inter-REM, while the wake activity decreased (NREM, P = 6.85e-7, R = 0.31; Wake, P = 0.006, R = −0.20, linear regression fit, n = 49 inter-REM intervals from n = 6 mice). Single dots represent the NREMs or wake activity of a mouse during the corresponding bin of a single inter-REM interval. Black line and shading, linear regression fit and 95% CI for the regression estimate. **(f)** Correlation between preceding REMs episode duration (REM_pre_) and mean NREMs or wake activity during the subsequent inter-REM interval. The average NREMs activity during inter-REM was negatively correlated with REM_pre_ (P = 0.029, R = −0.308, linear regression, n = 49 inter-REM intervals from n = 6 mice). There was no significant correlation between REM_pre_ and the subsequent wake activity during inter-REM (P = 0.846, R = −0.028). Each dot represents the activity from a mouse during a single inter-REM interval. Black line and shading, linear regression fit and 95% CI for the regression estimate.

